# Structure of LRRK1 and mechanisms of autoinhibition and activation

**DOI:** 10.1101/2022.11.22.517582

**Authors:** Janice M. Reimer, Andrea M. Dickey, Yu Xuan Lin, Robert G. Abrisch, Sebastian Mathea, Deep Chatterjee, Elizabeth J. Fay, Stefan Knapp, Matthew D. Daugherty, Samara L. Reck-Peterson, Andres E. Leschziner

**Author notes:** Correspondence should be addressed to S.L.R-P. or A.E.L. These authors contributed equally (Andrea M. Dickey, Yu Xuan Lin, Robert G. Abrisch).

## Abstract

Leucine Rich Repeat Kinase 1 and 2 (LRRK1 and LRRK2) are homologs in the ROCO family of proteins in humans. Despite their shared domain architecture and involvement in intracellular trafficking, their disease associations are strikingly different: LRRK2 is involved in familial Parkinson’s Disease (PD) while LRRK1 is linked to bone diseases. Furthermore, PD-linked mutations in LRRK2 are typically autosomal dominant gain-of-function while those in LRRK1 are autosomal recessive loss-of-function. To understand these differences, we solved cryo-EM structures of LRRK1 in its monomeric and dimeric forms. Both differ from the corresponding LRRK2 structures. Unlike LRRK2, which is sterically autoinhibited as a monomer, LRRK1 is sterically autoinhibited in a dimer-dependent manner. LRRK1 has an additional level of autoinhibition that prevents activation of the kinase and is absent in LRRK2. Finally, we place the structural signatures of LRRK1 and LRRK2 in the context of the evolution of the LRRK family of proteins.

## Introduction

ROCO proteins, discovered 20 years ago (Goldberg et al., 2002), are an unusual class of G-proteins distinguished by a Ras-like GTPase embedded in the context of a much larger polypeptide, in contrast to the more canonical G-proteins that function independently. This architecture led to the naming of these Ras-like domains as Ras-Of-Complex, or ROC. ROCO proteins are present in bacteria, archaea, plants, and metazoans (Wauters et al., 2019). All known ROCO proteins have an architectural domain immediately following the GTPase, termed a C-terminal Of Roc, or COR. This ROC-COR architecture is what gives rise to ROCO, the family name.

ROCO proteins in humans, of which there are four, gained prominence when mutations in one of its members, Leucine Rich Repeat Kinase 2 (LRRK2), were shown to be one of the most common causes of familial Parkinson’s Disease (PD) (Funayama et al., 2002; Simón-Sánchez et al., 2009; Zimprich et al., 2004). LRRK2 belongs to a class of ROCO proteins that, as its name indicates, contains a kinase domain in addition to its ROC GTPase, with the kinase immediately following the COR domain. Only one other ROCO protein in humans belongs to the same class: Leucine Rich Repeat Kinase 1 (LRRK1), LRRK2’s closest homolog. LRRK1 and LRRK2 have a very similar domain organization: an N-terminal half containing Ankyrin (ANK) and Leucine Rich Repeats (LRR), and a catalytic C-terminal half containing the ROC-COR ROCO signature, followed by the kinase and a WD40 domain (Figure 1A). The only difference between them is the presence of an Armadillo repeat domain at the N-terminus of LRRK2, which LRRK1 lacks. The similarities between LRRK1 and LRRK2 extend to their cell biological functions. Both proteins are involved in intracellular trafficking; they phosphorylate Rab GTPases that mark vesicular cargo that is transported along the microtubule cytoskeleton by the molecular motors dynein and kinesin (Malik et al., 2021; Steger et al., 2017, 2016). However, LRRK1 and LRRK2 phosphorylate non-overlapping sets of Rabs (Malik et al., 2021; Steger et al., 2017, 2016). For example, LRRK1 phosphorylates Rab7a, which is involved in the late endocytic pathway, while LRRK2 phosphorylates Rab8 and Rab10, which are involved in trans-Golgi transport (Pfeffer, 2017).

**Figure 1.**
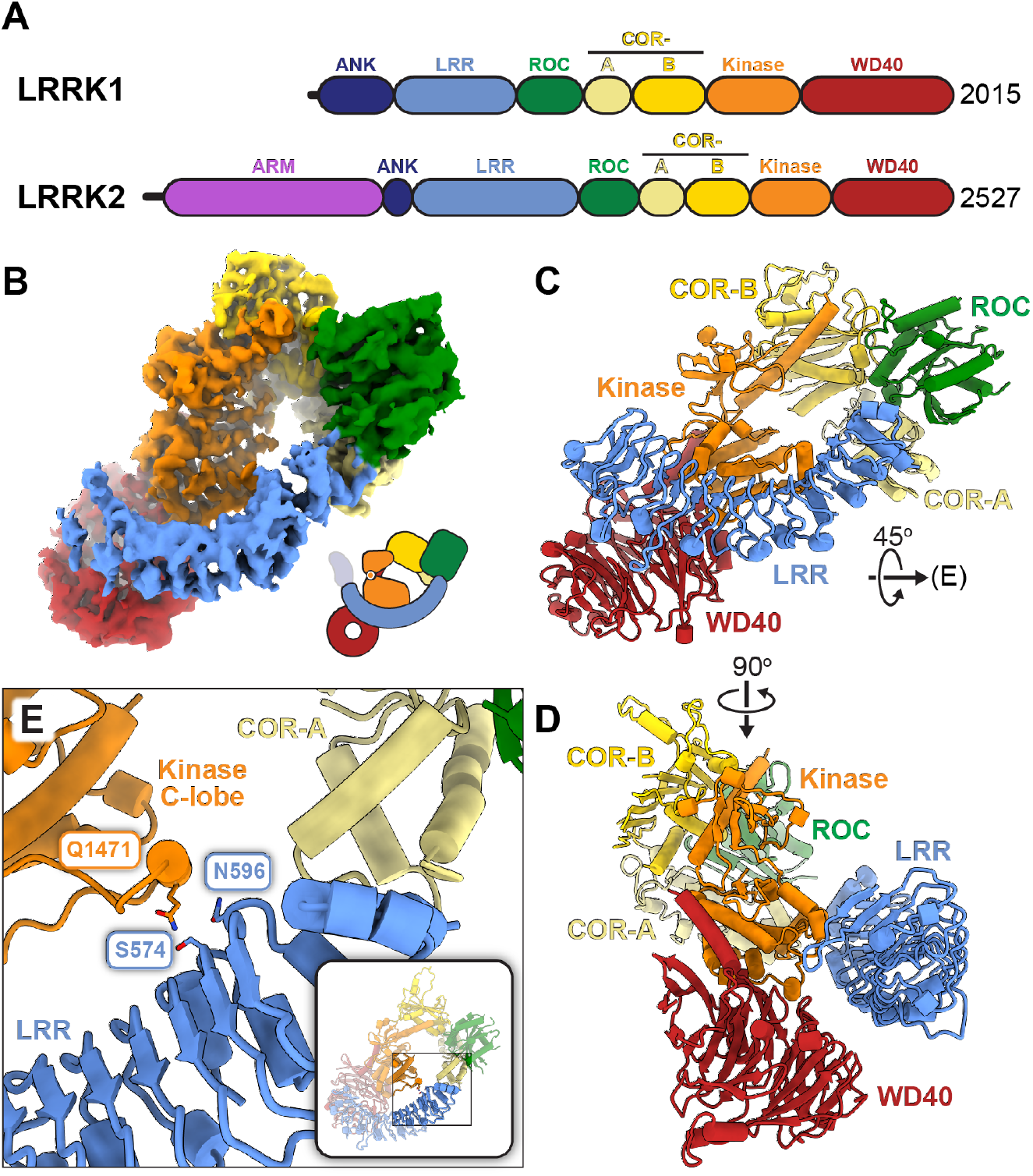
Cryo-EM structure of monomeric LRRK1. **(A)** Schematic domain organization of LRRK1 and LRRK2. The coloring scheme shown here is used in all figures. **(B-D)** Cryo-EM map (B) and two views of the model (C,D) of monomeric LRRK1. Note: The monomer model shown here is the one built after symmetry expansion of the LRRK1 dimer, as that map showed better density for the region highlighted in (E). The cartoon in (B) indicates that while present in our construct, the ANK domain is disordered in the cryo-EM map. **(E)** Close-up of the contact between the LRR repeat and the kinase’s C-lobe. The inset highlights the region of the model shown in the main panel.

Despite the molecular and cell biological similarities between LRRK1 and LRRK2, the proteins are strikingly different when it comes to their disease association. LRRK2 is well known for its involvement in Parkinson’s Disease; all the most common PD-linked mutations in LRRK2 are autosomal dominant gain-of-function mutations that activate its kinase (Ravinther et al., 2022). LRRK2 has also been linked to Crohn’s disease and leprosy (Hui et al., 2018; Schurr and Gros, 2009). In contrast, LRRK1 is not involved in PD and is instead linked to two rare bone diseases: osteopetrosis and osteosclerotic metaphyseal dysplasia (Xing et al., 2017). In further contrast with LRRK2, disease-linked mutations in LRRK1 are autosomal recessive, loss-of-function mutations (Xing et al., 2017). How two proteins with such similar domain architecture and related cellular functions are so different when it comes to their involvement in pathology remains a mystery.

While much is yet to be learned about LRRK2, our structural and mechanistic understanding of it has expanded significantly over the last few years, with several structures of the protein now available (Deniston et al., 2020; Myasnikov et al., 2021; Snead et al., 2022; Watanabe et al., 2020), along with insights into LRRK2’s cellular localization (Usmani et al., 2021), substrates (Malik et al., 2021; Steger et al., 2017, 2016), and regulation (Ravinther et al., 2022). This level of knowledge is missing for LRRK1. Understanding the differences and similarities between LRRK1 and LRRK2 would shed light into their unique cellular functions and how those lead to such different involvements in disease. Furthermore, determining what properties are LRRK1-specific would help us better define those that are unique to LRRK2, and thus likely to be involved in the etiology of LRRK2-associated PD.

We set out to bridge the gap in our understanding of LRRK1 by obtaining structures of full-length LRRK1 and comparing them with those of LRRK2. Here we report cryo-EM structures of full-length LRRK1 in its monomeric and dimeric forms. Despite LRRK1’s monomer having an overall structure similar to that of LRRK2, it differs from it in significant ways. LRRK2’s monomer is regulated by steric autoinhibition, with the LRR repeats blocking access to the kinase’s active site; this autoinhibition is unchanged in the inactive form of the dimer (Myasnikov et al., 2021). In contrast, monomeric LRRK1’s LRR repeats are shifted, making the kinase active site accessible. Surprisingly, LRRK1 achieves steric autoinhibition *in trans* by forming a dimer entirely unrelated to that formed by LRRK2; in it, the ANK repeats of each LRRK1 monomer block access to the kinase’s active site of the other monomer. LRRK1 also differs from LRRK2 in that its dimer is stabilized by a more complex set of interactions. Among them, we identified two interactions involving disordered regions of the protein that, when mutated, lead to increased phosphorylation of Rab7a, a LRRK1 substrate, in cells. Finally, we found that LRRK1 has a second level of autoinhibition, absent in LRRK2, where a loop arising from its COR-B domain reaches into the kinase’s active site and prevents the catalytic DYG motif from reaching its active conformation. This loop contains three sites that are targets for PKC phosphorylation (Malik et al., 2022). We show that mutations that destabilize the autoinhibitory conformation of the loop also increase phosphorylation of Rab7a in cells. Finally, we perform an evolutionary analysis of LRRK proteins to determine when characteristic structural features of LRRK1 and LRRK2 arose during metazoan evolution of the LRRK family of proteins.

## Results

### Cryo-EM structure of monomeric LRRK1

We purified LRRK1(Δ1-19), where the N-terminal 19 residues, predicted to be disordered, were deleted, and imaged it in the presence of Rab7a, ATP and GDP. Most particles classified as monomers, with a small subset forming dimers (Figure S1). The monomer particles yielded a 3.6Å structure of LRRK1(Δ1-19) (Figure 1B-D). Although the catalytic C-terminal half of LRRK1, which contains the ROC, COR, kinase, and WD40 domains (“RCKW”), adopts a Jshaped architecture similar to that of LRRK2 (Snead et al., 2022), the location of the N-terminal leucine rich repeat (LRR) domain differs between LRRK1 and LRRK2 in a functionally significant way. The N-terminal LRRs of LRRK2 are positioned in such a way that the LRR physically blocks access to the kinase’s active site (Figure 2A), in what appears to be an autoinhibited conformation (Myasnikov et al., 2021). In contrast, the LRRs of LRRK1 are shifted towards its WD40 domain, leaving its kinase’s active site exposed (Figures 1B, 2A). The only interaction made by the LRR domain, which extends from the GDP-bound ROC domain, with the rest of LRRK1(Δ1-19) is a small contact with the kinase’s C-lobe (Figure 1E). Interestingly, the residue in the kinase involved in this contact corresponds to N2081 in LRRK2, where a mutation linked to Crohn’s Disease has been identified (Hui et al., 2018). The kinase domain of LRRK1 is in the open, or inactive conformation with its DYG motif “out”, and even though the overall conformation of LRRK1(Δ1-19) would not prevent a Rab substrate from being engaged, we did not see any density for Rab7a in our LRRK1 map. This could be explained in part by the presence of an autoinhibitory loop extending from COR-B into the kinase active site, which we discuss in a separate section below. The ANK domain was either disordered or too flexible relative to the rest of the protein to be seen in our map.

**Figure 2.**
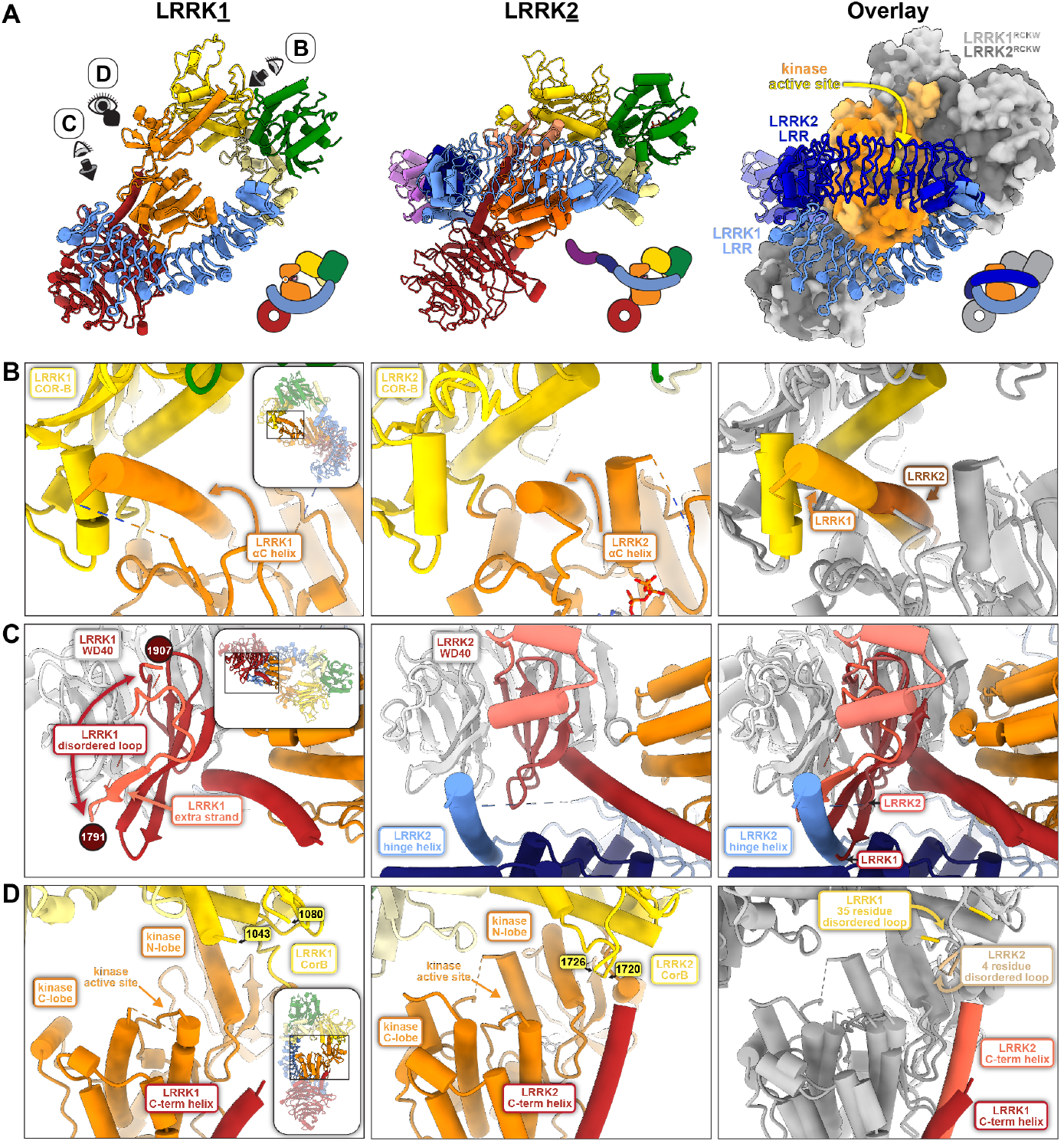
Comparison of monomeric LRRK1 and LRRK2. The LRR domain in LRRK1 does not sterically block access to the kinase’s active site as it does in LRRK2. Models and cartoons are shown for LRRK1 (left), LRRK2 (center), and an overlay of the two structures (right). In the overlay, the “RCKW” portion of LRRK1 and LRRK2 is shown as a surface representation, with the kinase in orange and the remaining domains in grey (light grey for LRRK1, dark grey for LRRK2). The directions of close-ups shown in panels (B-D) are indicated on the left-side panel. **(B-D)** These panels show comparisons between LRRK1 (left) and LRRK2 (center) focused on features that are different between the two structures, and a superposition to highlight those differences (right). The insets highlight the region of the structure shown in the main panel. (B) The αC helix in LRRK1’s kinase domain is several turns longer than its counterpart in LRRK2. (C) LRRK1’s WD40 domain has features that would clash with an element analogous to LRRK2’s latch helix. (D) LRRK1’s C-terminal helix is shorter than LRRK2’s.

An unusual feature of LRRK1 is the length of its kinase’s αC helix: it is ∼4 turns longer than is typical in kinases (Figure 2B). The extra residues pack tightly against the COR-B domain through an extensive hydrophobic interaction that is unique to LRRK1 (Figure S2A,B). Interestingly, the RCKW moiety of fulllength LRRK1 fits very well within a map of LRRK1_RCKW_ (Figure S2C), indicating that the presence of the N-terminal repeats does not alter the conformation of the catalytic half of the protein. This contrasts with LRRK2, where the position of the ROC-COR domains differs significantly between the full-length and LRRK2_RCKW_ structures (Figure S2D). It is possible that the more extensive interface between LRRK1’s αC helix and COR-B domain rigidifies the RCKW portion of LRRK1. Additionally, the longer helix is involved in one of LRRK1’s dimer interfaces, discussed below, which would not be possible with a helix of standard length.

Although LRRK1’s WD40 domain forms a typical seven-bladed beta propeller, the penultimate blade has several distinctive features (Figure 2C). Instead of the usual four strands, it starts with an additional strand, followed by a ∼112 residue disordered loop, unique to LRRK1, before the normal fold continues. The third and fourth strands are unusually long and would clash with where LRRK2’s hinge helix interacts with the WD40 domain (Figure 2C); LRRK1 lacks a hinge helix or an equivalent structural element that can interact with the WD40. LRRK2’s hinge helix is inserted at the start of the LRR domain and precedes a ∼100 residue disordered loop that contains key phosphorylation sites for binding members of the 14-3-3 family of proteins (Nichols et al., 2010), which are involved in regulating signaling in eukaryotic cells. The analogous, much shorter sequence in LRRK1 (residues 244-255) is disordered in our structure.

The C-terminal helix is a structural feature shared between the two LRRK proteins. In LRRK1, the last six residues of the protein are disordered, resulting in a C-terminal helix that is notably shorter than that of LRRK2 (Figure 2D). It was postulated that for LRRK2 the C-terminal helix, kinase N-lobe, and COR-B form a regulatory hub where phosphorylation of residue T2524 in the C-terminal helix could regulate kinase activity (Deniston et al., 2020). There are no known phosphorylation sites on LRRK1’s C-terminal helix and it does not extend far enough to contact the kinase Nlobe or COR-B. Since AlphaFold’s (Jumper et al., 2021) predicted LRRK1 structure has a fully folded C-terminal helix, we wanted to understand whether this is a result of AlphaFold’s LRRK1 being modeled in an active state or a more fundamental feature of unknown function. We deleted the last six residues of LRRK1 [LRRK1(Δ2010-2015)] and

### Cryo-EM structure of dimeric LRRK1

We purified full length LRRK1 and imaged it in the presence of GDP alone. Surprisingly, this construct, which contained the N-terminal 19 residues we had deleted in the construct we used for our monomer structure, yielded almost exclusively dimers (Figure S4). The sample still showed severe preferred orientation, but we were able to partially overcome this with the addition of the detergent brij-35 and obtained a 4.6 Å structure of the full-length LRRK1 dimer (Figure 3). Each molecule in the dimer has the same conformation we observed in the monomer; however, the ANK domain is fully resolved in the dimer structure.

**Figure 3.**
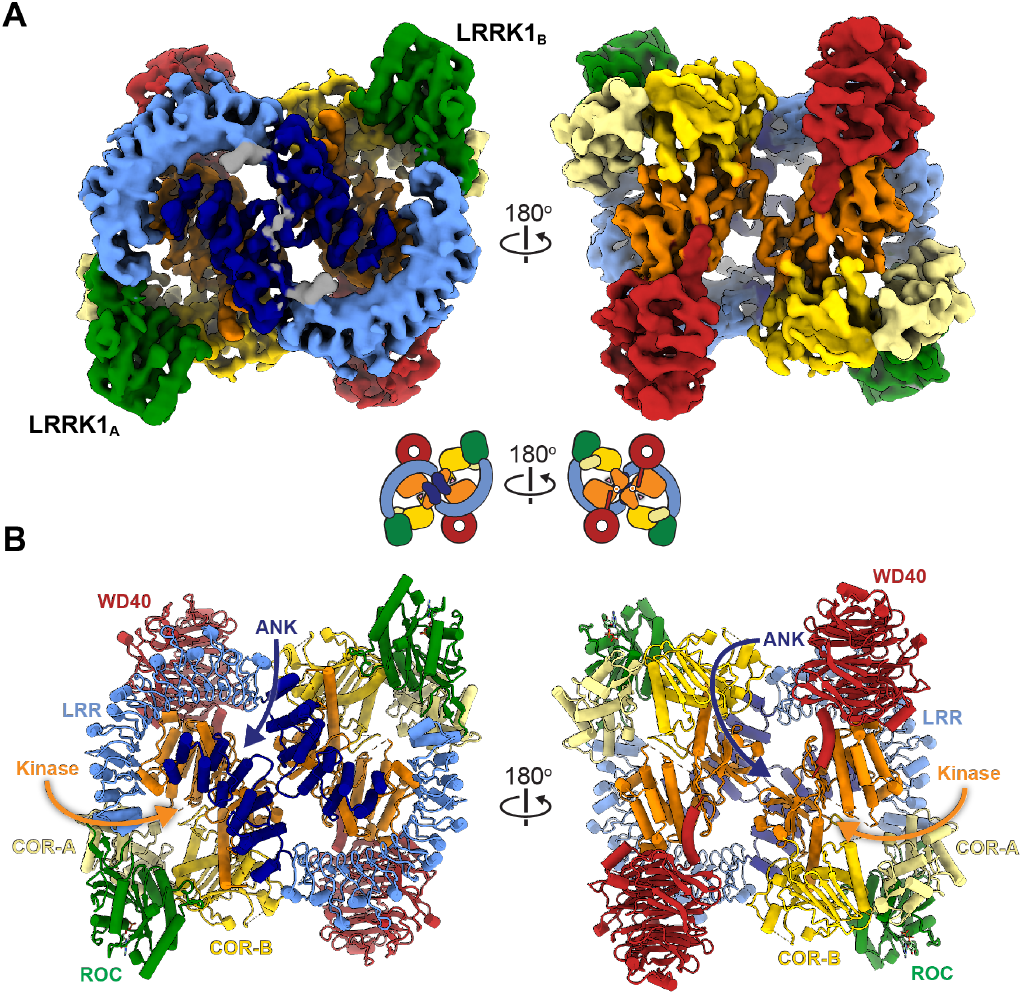
Cryo-EM structure of dimeric LRRK1. **(A, B)** Cryo-EM map (A) and model (B) of dimeric LRRK1. Domains are indicated for the LRRK1A monomer in (B). measured phosphorylation of Rab7a, a LRRK1 substrate (Malik et al., 2021), in cells. We did not observe a significant difference in Rab7a phosphorylation between full-length LRRK1 and LRRK1(Δ2010-2015) (Figure S3), suggesting that the end of the C-terminal helix does not play a major role in LRRK1 regulation.

Surprisingly, despite the similarities in their domain organization and the structures of the monomers, the LRRK1 dimer bears no resemblance to that of LRRK2 (Figure 4A,B). While LRRK2 forms a parallel dimer mediated by a single homotypic interaction involving its COR-B domain (Figure 4B), LRRK1 forms an antiparallel dimer mediated by several homo- and heterotypic interactions (Figure 4A). The ANKLRR domains of each LRRK1 wrap around those of the other, making symmetrical contacts between the ANK:ANK and LRR:ANK domains (Figure 4C,D). While the resolution is too low to determine specific interactions at the ANK:LRR interface, the surfaces involved are electrostatically complementary, lined with acidic residues in the ANK domain and basic residues in the LRR domain (Figure 4D). A major interaction interface is formed by the kinase C-lobe and LRR domains of one monomer and the opposite molecule’s ANK domain (Figure 4E). As mentioned above, the C-lobe contact site is analogous to the site in LRRK2 linked to Crohn’s disease (N2081D). In contrast to LRRK2, whose dimer is mediated entirely by the COR-B domain in its RCKW moiety, the LRRK1 dimer shows minimal direct interactions between RCKWs; the only contact seen in our structure is a homotypic interaction involving residues 1263-1265 in the kinase N-lobe (Figure 4F).

**Figure 4.**
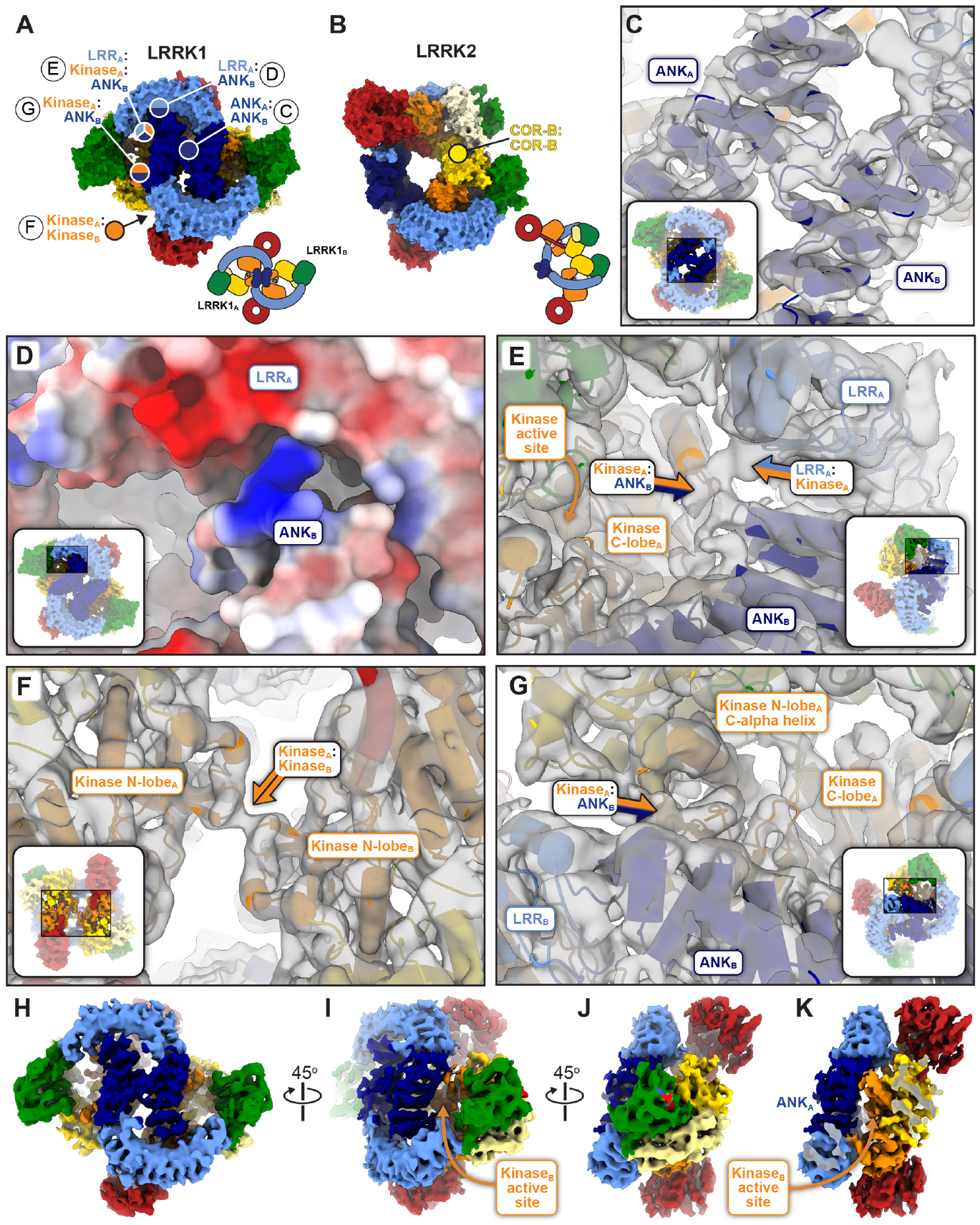
Comparison of dimeric LRRK1 and LRRK2. **(A**,**B)** Models of LRRK1 (A) and LRRK2 (B) shown in surface representation. The models are shown with the bottom-right monomer in the same orientation to highlight the differences in the architecture of the dimer. The different interfaces involved in forming the LRRK1 (A) and LRRK2 dimers are indicated. In the case of LRRK1, panel letters next to the interface labels refer to the detailed views shown in this figure. Cartoons of the dimers are shown below the models. **(C-G)** Close ups of the different LRRK1 dimer interfaces. The insets highlight the area of the structure shown in the main panel. Except for (D), all other panels show the LRRK1 model inside the cryo-EM map. (C) Symmetric interface formed by the ANK domains. (D) The interface formed by the ANK domain of one monomer and the LRR domain of the other monomer brings together surfaces of complementary charge. (E) The kinase C-lobe of one monomer interacts both with the LRR domain of the same monomer and the ANK domain of the other monomer. Along with the interaction shown in (G), this anchors the ANK domain on top of the kinase, where it blocks access to the active site. (F) Symmetric interaction between the N-lobes of the kinases. This is the only dimeric interface involving only a domain in the C-terminal half of LRRK1 (RCKW). (G) The N-lobe of the kinase of one monomer interacts with the ANK domain of the other monomer. Along with the interaction shown in (E), this anchors the ANK domain on top of the kinase, where it blocks access to the active site. **(H)** The Cryo-EM map of the LRRK1 dimer, colored by domains, is shown in the same orientation as the model in (A). **(I)** Rotated view of the dimer map showing how the kinase active site is buried. **(J, K)** An additional rotation of the dimer (J) and clipping of the density in front (K) highlights how the kinase from one monomer is buried under the ANK domain from the other monomer.

### LRRK1 is sterically autoinhibited in trans in the dimer

The main contact between the two LRRK1 monomers, and the most striking feature of the dimer, involves the ANK domains; each ANK domain contacts the opposite molecule’s kinase on both its N- and C-lobes (Figure 4H-K), effectively blocking access to the kinase. Our structure suggests that the LRRK1 dimer uses the ANK domain to accomplish, *in trans*, the type of steric autoinhibition achieved by the LRR domain in the LRRK2 monomer.

### LRRK1’s N-terminus stabilizes the dimer

As mentioned above, the LRRK1 (full-length) and LRRK1(Δ1-19) constructs showed dramatically different distribution of monomers and dimers on our cryo-EM grids despite differing only in the presence or absence of 19 residues at the N-terminus; full-length LRRK1 resulted almost exclusively in dimers, while the LRRK1(Δ1-19) truncated construct was mainly monomeric. This was particularly surprising given that AlphaFold predicts that the first ∼48 residues of LRRK1 are disordered (Figure 5A), and the fact that the first residue we were able to model in the LRRK1 dimer was R51 (Figure 5B). However, we noted two areas of density in our cryo-EM map, near the junction of the ANK-LRR domains and on top of the ANK:ANK interaction, that were not accounted for by our or the AlphaFold models (Figure 5C). We hypothesized that these densities could be accounted for by the extreme N-terminus of LRRK1 docking onto the ANK-LRR to stabilize the autoinhibited dimer. A prediction from this hypothesis was that deleting the N-terminus should destabilize the dimer and thus increase LRRK1 kinase activity. We tested this by measuring phosphorylation of Rab7a in cells. We expressed one of three constructs in 293T cells: full-length LRRK1, a 25-residue N-terminal deletion [LRRK1(Δ1-25)], and a 48-residue deletion [LRRK1(Δ1-48)], based on AlphaFold’s prediction that the first 48 residues in LRRK1 would be disordered. In agreement with our hypothesis that the N-terminus plays a role in stabilizing the autoinhibited dimer, both deletions resulted in a ∼50% increase in phosphorylation of Rab7a compared to full-length LRRK1 (Figure 5D). This is comparable to what we observed with the hyperactive kinase mutant LRRK1_K746G_ (equivalent to the Parkinson’s Disease-linked R1441G mutation in LRRK2) (Figure 5D). The difference in activity between LRRK1(Δ1-25) and LRRK1(Δ1-48) was not statistically significant, suggesting that the first 25 residues are involved in stabilizing the LRRK1 dimer.

**Figure 5.**
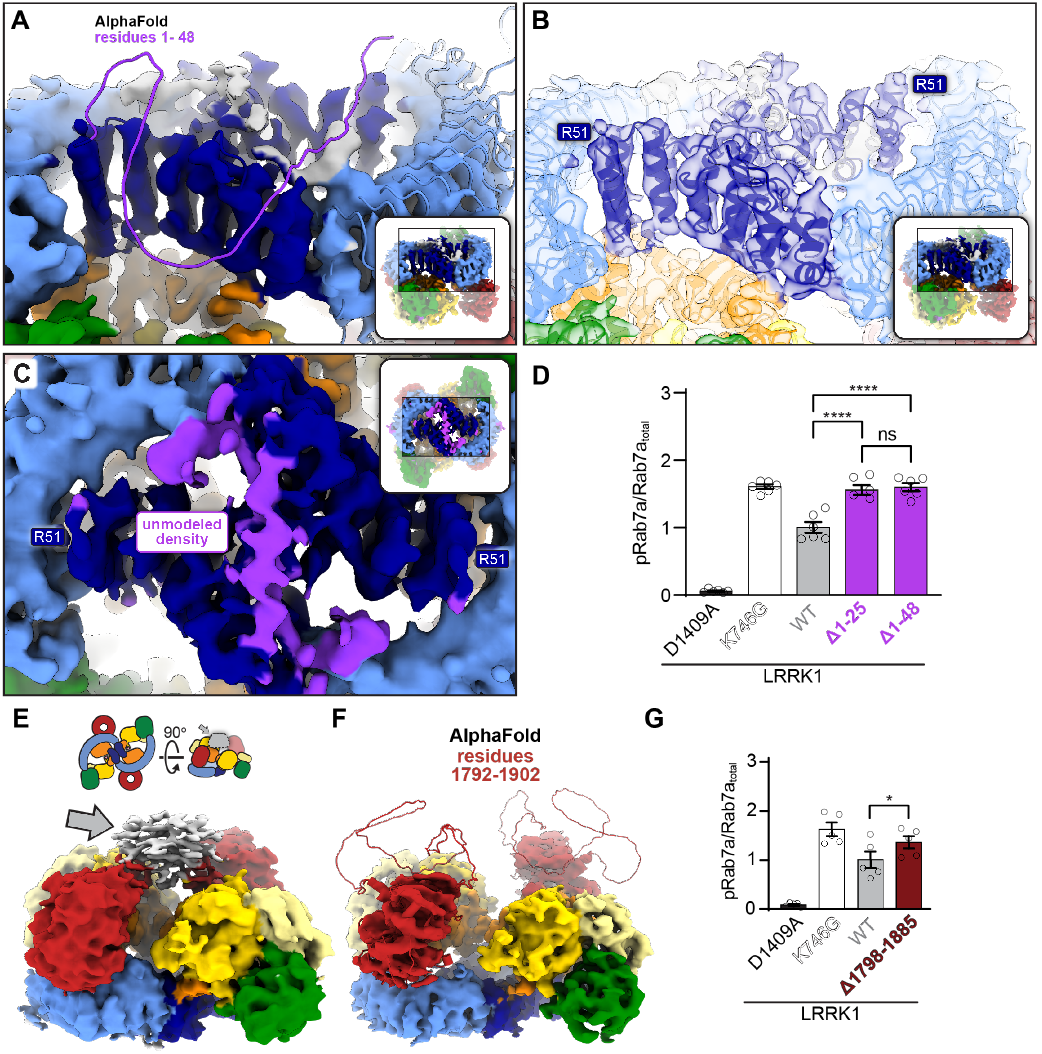
Disordered loops in LRRK1’s ANK and WD40 domains help stabilize the autoinhibited dimer. **(A)** The ANK and LRR domains of the AlphaFold model of human LRRK1 (Q38SD2) are shown docked into our cryo-EM map of LRRK1’s dimer. The N-terminal residues 1-48, which are unstructured in the AlphaFold model and not included in our model, are shown in purple. The insets in panels (A-C) highlight the area of the structure shown in the main panel. **(B)** Same view as in (A) with our model of the LRRK1 dimer shown inside the cryo-EM map to highlight that R51 is the first residue that was modeled in our structure. **(C)** View of the ANK-ANK interface in the cryo-EM map of the LRRK1 dimer. The purple density corresponds to the region of the cryo-EM map unaccounted for by our current model. The location of the residues where our model begins (R51) are indicated. **(D)** Rab7a phosphorylation in 293T cells expressing full-length LRRK1 or N-terminally truncated constructs missing the first 25 or 48 residues. LRRK1(K746G), which is known to increase Rab7 phosphorylation in cells, and LRRK1(D1409A), which is known to be kinase inactive, were tested as well. 293T cells were transiently transfected with the indicated plasmids encoding FLAG-LRRK1 (wild type or mutant) and GFP-Rab7. Thirty-six hours post-transfection the cells were lysed, immunoblotted for phosphor-Rab7 (pS72), total GFP-Rab7, and total LRRK1. The mean ± s.e.m. is shown, ****p<0.0001, ns=not significant, one-way ANOVA. Individual data points represent separate populations of cells obtained across at least three independent experiments (n ≥ 3). **(E)** We observed weak density (grey arrow) connecting the WD40 domains in our initial dimer maps. The cartoons above the map indicate the orientation of the maps shown in (E) and (F) as well as the location of the weak density. **(F)** The final cryo-EM map of the LRRK1 dimer with the WD40 domain from the AlphaFold model of LRRK1 docked in; AlphaFold predicted that residues 1792-1902 in the WD40 domain form an extended disordered loop (shown). **(G)** Rab7a phosphorylation in 293T cells expressing GFP-Rab7a and full-length WT LRRK1 or a LRRK1 variant missing residues 1798-1885 from its WD40 domain. LRRK1(K746G), which is known to increase Rab7 phosphorylation in cells, and LRRK1(D1409A), which is known to be kinase inactive, were tested as well. 293T cells were transiently transfected with the indicated plasmids encoding FLAG-LRRK1 (wild type or mutant) and GFP-Rab7. Thirty-six hours post-transfection the cells were lysed, immunoblotted for phosphor-Rab7 (pS72), total GFP-Rab7, and total LRRK1. The mean ± s.e.m. is shown. *p=0.0317, one-way ANOVA Individual data points represent separate populations of cells obtained across at least three independent experiments (n ≥ 3).

### LRRK1’s dimer is also stabilized by a loop in the WD40 domain

While processing data for the LRRK1 dimer, we noticed a large density, adjacent to the C-terminal helices, connecting the two WD40 domains (Figure 5E); this density was weak and noisy and was only seen when the map was displayed at lower threshold. The mask used in processing was autogenerated in cryoSPARC (Punjani et al., 2017a), and we first wondered if this was an artifact due either to dynamic masking during processing or to the two-fold symmetry applied to the map of the dimer. To test this, we reprocessed the data either using a mask that excluded the region where the density had appeared, or without applying symmetry. In both cases, the unaccounted-for density persisted. We next wondered if the long LRRK1-specific loop in the WD40 domain (residues 1791-1907, Figure 5F), which we had not been able to model, could be involved in forming the large density. We hypothesized that this loop might stabilize the autoinhibited LRRK1 dimer, as was the case for the N-terminus, and thus that its removal would increase LRRK1’s kinase activity. We engineered a deletion of most of this loop, LRRK1(Δ1798-1885), that was predicted (by AlphaFold modeling) to maintain proper folding of the WD40 domain. As we had done before, we measured phosphorylation of Rab7a in 293T cells expressing either full-length LRRK1, or the LRRK1(Δ1798-1885) construct. In agreement with our hypothesis, deletion of the WD40 loop resulted in a significant increase in Rab7a phosphorylation in cells (Figure 5G).

### A loop from COR-B inhibits LRRK1’s kinase

Our initial model for the kinase domain of LRRK1 showed that the DYG motif, a tripeptide involved in ATP binding, was in the “out”, or inactive conformation, but it also revealed additional density in the back pocket of the kinase that was not accounted for by the model. The density was located where Y1410 from the DYG motif would dock in the DYG “in”, or active conformation. Further processing of the dimer dataset using symmetry expansion and focused refinement provided a surprising explanation for this unaccounted density, which is present in both the monomer and dimer maps. A loop from the COR-B domain (residues 1048-1082), which is predicted by AlphaFold to be entirely disordered (Figure S5A, B), threads into the kinase domain active site (Figure 6A); approximately half of the COR-B loop is ordered in our map. The equivalent loop in LRRK2 is half as long and does not extend towards the active site (Figure S5C). The side chain of F1065, at the tip of the loop, sits inside the back pocket of the kinase (Figure 6B), occupying the position of Y1410 in the DYG-in conformation. A similar “plugging” of the kinase back pocket was observed in the DDR1 kinase (Figure S5D-F) (Sammon et al., 2020). This suggests that the COR-B loop is an autoinhibitory element in LRRK1. To test this idea, we measured Rab7a phosphorylation in 293T cells expressing either wild-type LRRK1 or LRRK1(F1065A), which we expected would at least partly relieve the autoinhibition. In agreement with this, the F1065A mutation led to a two-fold increase in the level of Rab7a phosphorylation in cells, comparable to that observed with the hyperactive K746G mutant (Figure 6C).

**Figure 6.**
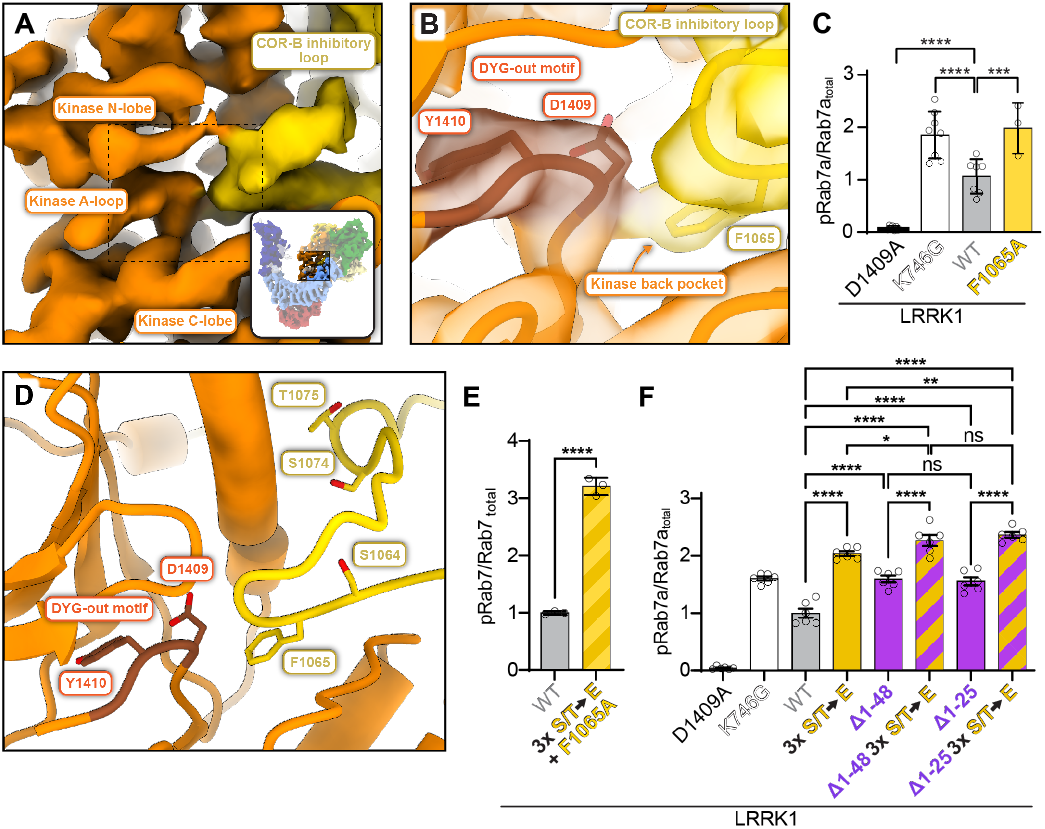
A loop from the COR-B domain directly inhibits LRRK1’s kinase. Close up of the kinase domain in the cryo-EM map of monomeric LRRK1; the density colored in yellow corresponds to a loop arising from the COR-B domain that reaches towards the kinase active site. The inset highlights the area of the structure shown in the main panel. The dashed outline indicates the region shown in panel (B). **(B)** Our model of LRRK1 is shown inside the cryo-EM map around the kinase’s active site. The DYG motif, in its “out” conformation, is shown. This panel also shows that F1065, a residue in the COR-B inhibitory loop, occupies the kinase’s “back pocket”, where Y1410 must dock to bring the DYG motif into its “in”, or active conformation. **(C)** Rab7a phosphorylation in cells expressing full-length LRRK1 WT or carrying a F1065A mutation. LRRK1(K746G), which is known to increase Rab7 phosphorylation in cells, and LRRK1(D1409A), which is known to be kinase inactive, were tested as well. 293T cells were transiently transfected with the indicated plasmids encoding for FLAG-LRRK1 (wild type or mutant) and GFP-Rab7. Thirty-six hours post-transfection the cells were lysed, immunoblotted for phosphor-Rab7 (pS72), total GFP-Rab7, and total LRRK1. The mean ± s.e.m. is shown. ****p<0.0001, ***p=0.001, one-way ANOVA. Individual data points represent separate populations of cells obtained across at least three independent experiments (n ≥ 3). **(D)** An expanded view of the area shown in (B), without the cryo-EM map. The three sites of PKC phosphorylation in the COR-B inhibitory loop—S1064, S1074, and S1075—are shown in addition to F1065. **(E, F)** Rab7a phosphorylation in 293T cells expressing GFP-Rab7a and full-length WT LRRK1 or LRRK1 carrying a combination of the F1065A mutation with three phosphomimetic mutations (S1064E/S1074E/T1075E) in the COR-B inhibitory loop (E) or truncated (Δ1-48 and Δ1-25) versions of LRRK1 with or without the phosphomimetic mutations (S1064E/S1074E/T1075E) in the COR-B inhibitory loop (F). The triple phosphomimetic mutant is abbreviated as “S/T → E” in the graphs. LRRK1(K746G), which is known to increase Rab7 phosphorylation in cells, and LRRK1(D1409A), which is known to be kinase inactive, were tested as well. 293T cells were transiently transfected with the indicated plasmids encoding for FLAG-LRRK1 (wild type or mutant) and GFP-Rab7. Thirty-six hours post-transfection the cells were lysed, immunoblotted for phosphor-Rab7 (pS72), total GFP-Rab7, and total LRRK1. The mean ± s.e.m. is shown. ****p<0.0001, **p,0.0021, *p,0.0032, ns=not significant. one-way ANOVA. Individual data points represent separate populations of cells obtained across at least three independent experiments (n ≥ 3).

### Phosphorylation of residues in the COR-B loop relieves LRRK1 autoinhibition

LRRK1 contains several consensus sites for phosphorylation by Protein Kinase C (PKC) (Malik et al., 2022). Three of these sites—S1064, T1074 and S1075—are found in the autoinhibitory COR-B loop (Figure 6D). Phosphorylation of these residues significantly increases the kinase activity of LRRK1 (Malik et al., 2022). In addition, preventing phosphorylation (by mutating the residues to alanine) reduces Rab7a phosphorylation in cells, while phosphomimetic mutations (to glutamate) increase it, although to a lesser extent than phosphorylation (Malik et al., 2022). Our structure provides a mechanistic explanation for this activation: phosphorylation of these residues disrupts the loop, which is nestled against the kinase domain, releasing F1065 from the kinase’s back pocket and allowing the DYG motif to adopt the active “in” conformation. Based on this model, we would expect some synergy between the F1065A mutation and phosphomimetic mutations in the COR-B loop (S1064E/T1074E/S1075). In agreement with this, LRRK1 carrying all four mutations resulted in a 3-fold increase in Rab7a phosphorylation in cells (Figure 6E).

Our work revealed two separate autoinhibitory mechanisms in LRRK1: (1) autoinhibition by the COR-B loop, which is present in both the monomer and dimer structures, and (2) steric autoinhibition of the kinase by the ANK domain, which occurs in trans and is dependent on dimerization. Given the seemingly independent nature of these mechanisms, we wondered if their effects would be additive to any extent. To test this, we introduced the phosphomimetic mutations in the context of the N-terminal deletions we had tested earlier: LRRK1(Δ1-25)(S1064E/S1074E/T1075E) and LRRK1(Δ1-48)(S1064E/S1074E/T1075E). As shown previously (Malik et al., 2022), Rab7a phosphorylation in 293T cells expressing full-length LRRK1 carrying the triple phosphomimetic mutations increased by a factor of 2 relative to wild-type LRRK1 (Figure 6F). Combining these mutations with either the 1-25 or 1-48 N-terminal truncation of LRRK1 resulted in a small, but statistically significant additional increase in Rab7a phosphorylation in cells (Figure 6F), suggesting that the effects from these two autoinhibitory mechanisms are additive.

### An evolutionary analysis of the structural signatures of LRRK1 and LRRK2

Structural information on LRRK2 has built up over the last few years (Deniston et al., 2020; Myasnikov et al., 2021; Snead et al., 2022; Watanabe et al., 2020). The data we presented here on LRRK1 now allow us to establish the structural signatures that define these two proteins. We set out to analyze the conservation of these features throughout evolution to understand which ones are most likely to be tied to LRRK1-or LRRK2-specific biological functions. We expect that this information will shed light on the etiology of PD and bone diseases.

We began by evaluating the evolutionary origin and phylogenetic distribution of LRRK proteins, defined as those with 40% or higher sequence coverage relative to human LRRK1 or LRRK2, to ensure complete coverage of the ROC, COR-A, COR-B, and kinase domains (see Methods). We found that LRRK proteins are present in a wide range of metazoan species, and that related proteins are present in amoeba. Phylogenetic analyses of these proteins revealed five distinct and well-supported LRRK clades, with amoeba proteins forming a single clade and the remaining four clades containing only metazoan proteins (Figure 7A and Figure S6), as has been observed previously (Marín, 2008). Human LRRK1 and LRRK2 are found in distinct clades that contain proteins from vertebrates, echinoderms (*e*.*g*., starfish), spiralians (*e*.*g*., mollusks), and cnidarians (*e*.*g*., corals). Arthropod and nematode LRRK proteins, which are annotated as either LRRK1 or LRRK2, are in fact found in a clade (labeled LRRK3 in Figure 7A) that is distinct from vertebrate LRRK1 and LRRK2, and that also contains proteins from echinoderms, spiralians, and cnidarians. Finally, a fourth clade (labeled LRRK4) contains only proteins from cnidarians. These data suggest, as has been proposed earlier (Marín, 2008), that gene duplication early in metazoan evolution led to four distinct LRRK protein families, followed by loss of individual LRRK members in different lineages. Notably, while vertebrates have only retained LRRK1 and LRRK2, arthropods and nematodes have retained only LRRK3, while spiralians and echinoderms have retained LRRK1, LRRK2, and LRRK3, and cnidarians have proteins from all four LRRK protein families (Figure 7B).

**Figure 7.**
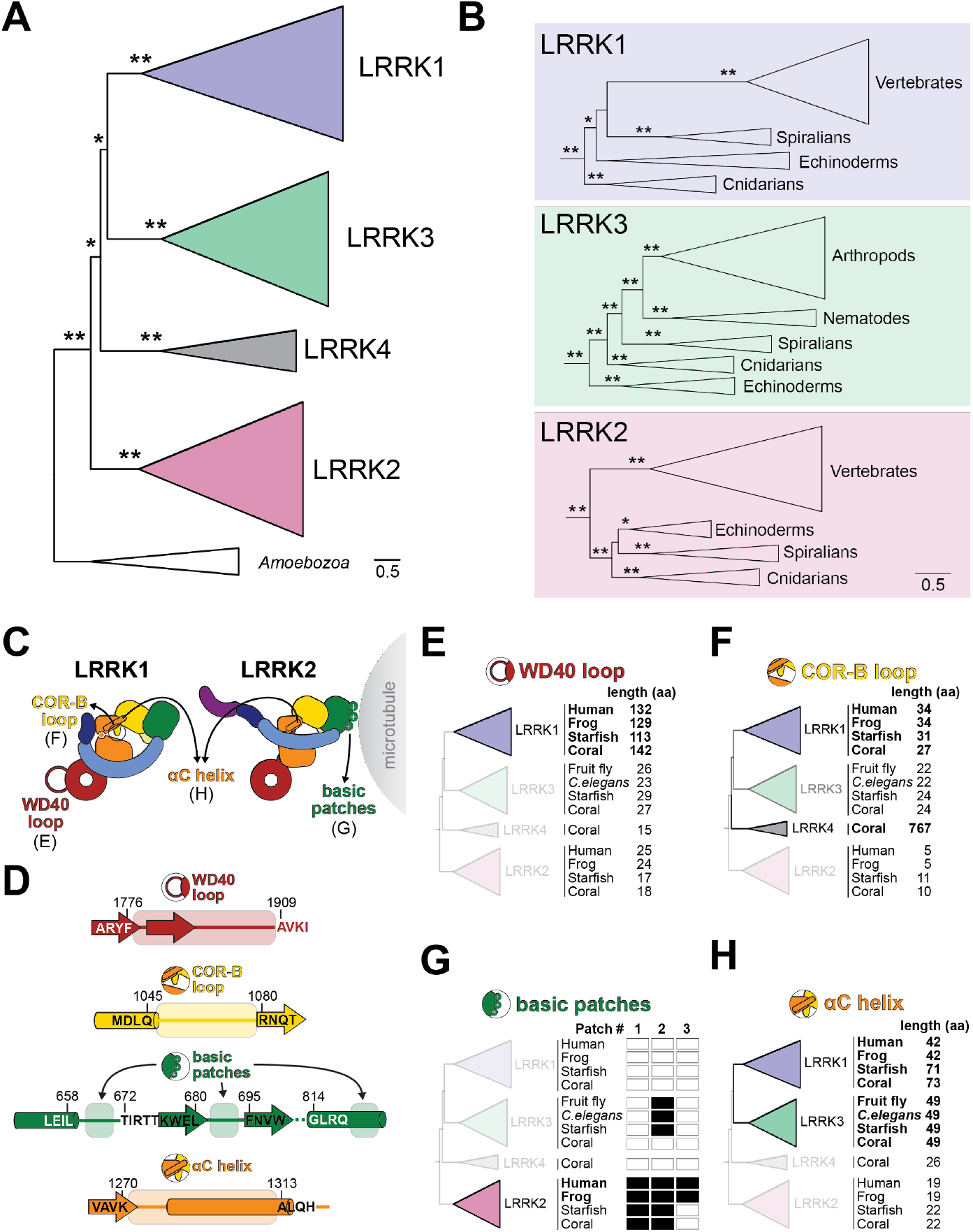
Evolution of structural motifs in LRRK proteins. Maximum likelihood phylogenetic tree of LRRK protein homologs (see supplemental datasets 1, 2, and 3 for complete listing of proteins, protein alignment, and complete phylogenetic tree, respectively). Shaded clades are LRRK proteins found in metazoans, using nomenclature proposed in (Marín, 2008). The tree is rooted on the only non-metazoan LRRK homologs, which are found in *Amoebozoa*, but the validity of those proteins as the ancestor to metazoan LRRK proteins is not well-established (Marín et al., 2008). Asterisks indicate bootstrap branch support (* >75% support, ** 100% support). **(B)** Expanded views of the phylogenetic tree shown in (A), highlighting major metazoan clades that contain members of LRRK1, LRRK2 and LRRK3. Asterisks indicate bootstrap branch support as in panel A. **(C)** Cartoons of LRRK1 and LRRK2 showing structural features whose conservation was analyzed here. The panels where results are presented for each feature are indicated. **(D)** Schematic of the boundaries used for determining whether one of the features shown in (C) is present in a given LRRK. Amino acid sequences and numbers are shown for regions flanking the sequences whose lengths (panels E, F, and H) or where the presence of key residues in the COR-B autoinhibitory loop (panel F), or basic patches (panel G) were measured in our analysis. **(E)** Representative proteins from each metazoan LRRK protein family were sampled from vertebrates (human: *Homo sapiens* and frog: *Xenopus laevis*), echinoderms (starfish: *Asterias rubens*), cnidarians (coral: *Dendronephthya giganteas*), arthropods (fruit fly: *Drosophila sechellia*) and nematodes (*Caenorhabditis elegans*). The length of the WD40 loop, as measured between the well-aligning motifs shown in (D), is shown next to each homolog. **(F)** As in panel E, except measuring the length of the CORB loop, as measured between the well-aligning motifs shown in (D). Filled boxes indicate the presence of key residues involved in autoinhibition and activation—the Phe that docks into the kinase back pocket (“F”), and the three phosphorylation sites (“P1-P3”). **(G)** As in panel E, except querying for the presence of basic patches. Filled boxes indicate the presence of a basic patch as defined by three basic amino acids in a stretch of four residues between the regions defined in (D). **(H)** As in panel E, except measuring the length of the region containing the αC helix, as measured between the well-aligning motifs shown in (D).

Using this LRRK protein phylogeny, we next asked when characteristic features of LRRK1 and LRRK2 arose during metazoan LRRK protein family evolution. The major structural features of LRRK2 that distinguish it from LRRK1 include: (1) the presence of basic residues in its ROC domain that allow it to bind to microtubules *in vitro* (Deniston et al., 2020; Snead et al., 2022) and, under some conditions, in cells (Berger et al., 2010; Bonet-Ponce et al., 2020; Deniston et al., 2020; Eguchi et al., 2018; Gomez et al., 2019; Purlyte et al., 2018; Snead et al., 2022); (2) the COR-B:COR-B interface that mediates dimerization (Myasnikov et al., 2021); (3) the two interfaces involved in formation of the microtubule-associated filaments: the same COR-B:COR-B interface that forms the dimer (Snead et al., 2022) and a WD40:WD40 interaction (Snead et al., 2022; Watanabe et al., 2020; Zhang et al., 2019) ; and (4) its long C-terminal helix that emerges from the WD40 domain and runs along the back of the kinase domain (Deniston et al., 2020). LRRK1-specific features include: (1) its disordered N-terminus, involved in stabilizing the autoinhibited dimer (Figure 5A-D); (2) the LRRK1-specific WD40 loop, also involved in stabilizing the autoinhibited dimer (Figure 5E-G); (3) the LRRK1-specific autoinhibitory COR-B loop (Figure 6); (4) the unusually long αC helix in its kinase domain (Figure 2B); and (5) a C-terminal helix that is shorter than the equivalent one in LRRK2 and makes fewer contacts with the kinase domain (Figure 2D). We focused our evolutionary analysis on four of these structural features: (1) the LRRK1-specific WD40 loop involved in autoinhibition, (2) the LRRK1-specific COR-B loop involved in autoinhibition, (3) the LRRK2-specific basic patches found in the ROC domain that mediate microtubule binding, and (4) the length of the αC helix in the kinase’s N-lobe, which is significantly longer in LRRK1 compared to LRRK2 (Figure 7C, D). We were not able to analyze two other features—the presence of the COR-B:COR-B and WD40:WD40 dimerization interfaces in LRRK2 that are required for the formation of the microtubule-associated filaments, and the differences in length in the C-terminal helix that emerges from the WD40 domain —due to the fact that the protein alignment in these regions was not of sufficient quality to confidently infer relatedness.

Consistent with the role that the WD40 loop plays in LRRK1 regulation, we found that this is a conserved feature of metazoan LRRK1s but is not found in any other metazoan LRRKs (Figure 7D, E). We also found that an extended (>30 residue) COR-B loop is conserved in LRRK1 proteins from vertebrates, echinoderms, and spiralians, although the regulatory elements in the loop—the Phe that occupies the kinase back pocket (F1065 in human LRRK1), and the three PKC phosphorylation sites (S1064/S1074/T1075 in human LRRK1) are only present in LRRK1s from jawed vertebrates (Figure 7D, F, Figure S7). Cnidarian LRRK1s have a short (<10 residue) loop in COR-B that resembles the length of the loop in all LRRK2s. Our structure of LRRK1 suggests that the COR-B loop, as defined in Figure 7D, must be at least 26 residues long for the autoinhibitory mechanism we identified to be possible. Interestingly, LRRK3s, including those from arthropods and nematodes, have an intermediate length loop, ranging from 20-30 residues, whereas most LRRK4s have a much longer insert (200 or more residues) in the COR-B domain (Figure 7F). The functional consequences of these differences in COR-B remain to be determined. Next, we examined metazoan LRRK proteins for the presence of the basic patches found in the LRRK2 ROC domain (Snead et al., 2022). The first two basic patches are found in loop regions while the third is found in an alpha helix (Figure 7D). Only in jawed vertebrate LRRK2s do all three basic patches exist in the analogous positions in the ROC domain (Figure 7G). In the jawless sea lamprey and more distantly related non-vertebrate metazoans, the patch 3 motif is not basic (Figure S8). Given that mutations in any of the three basic patches in LRRK2’s ROC domain significantly reduce microtubule binding in cells (Snead et al., 2022), our data suggest that LRRK2 from jawed vertebrates may be the only LRRKs that exhibit this property. Finally, we looked at the length of the region encompassing the kinase’s αC helix in the different LRRK proteins (Figure 7D). This region is longer in LRRK1s in vertebrates, echinoderms, and spiralians, and in LRRK3s in arthropods, nematodes, echinoderms, and spiralians, but shorter in LRRK2s and LRRK4s (Figure 7H). While our structure of LRRK1 showed an unusually long αC helix, it remains to be determined whether this is a feature of all LRRKs containing a longer insert in this part of the kinase.

## Discussion

Our structural and functional analysis of LRRK1 has revealed that although LRRK1 and LRRK2 have a similar domain architecture and related kinase substrates, the two proteins differ in the structures of their monomers and dimers and, most dramatically, in their mechanisms of autoinhibition and activation.

### LRRK1 regulation

One of the most striking aspects of our findings is the extent to which LRRK1 appears to be more stringently autoinhibited than LRRK2. Our current understanding of LRRK2 suggests that its activation should only require movement of the N-terminal repeats to relieve the physical blockage of the kinase’s active site by the LRR. Structures and structure predictions of LRRK2 indicate that this activation could take place in the context of either a monomer or a dimer as dimerization does not bring about new properties, at least in the context of the inactive dimer form (Myasnikov et al., 2021). In contrast, the equivalent steric inhibition in LRRK1, mediated by its ANK repeats, is dependent on dimerization. Given that the ANK repeats are also involved in mediating dimerization, along with several other homotypic and heterotypic interactions in LRRK1, it is not easy to see how autoinhibition in LRRK1 could be relieved without disrupting the dimer. Our work revealed two interactions involved in stabilizing the dimer: one mediated by the extreme N-terminus of LRRK1, and the other by the LRRK1-specific WD40 loops. Although disruption of either interface resulted in increased Rab7a phosphorylation in cells, this increase was relatively modest, suggesting that other interfaces still play a major role in stabilizing the LRRK1 dimer. How could this dimerization be regulated in cells? An intriguing observation is the presence of multiple predicted phosphorylation sites (NetPhos–3.1 server: https://services.healthtech.dtu.dk/service.php?NetPhos-3.1) in the two regions we identified as mediators of dimerization. Future studies will determine whether phosphorylation of some of these residues disrupts, or stabilizes, the autoinhibited LRRK1 dimer.

The autoinhibitory COR-B loop is unique to LRRK1 and provides an additional level of regulation beyond that brought about by dimerization; this loop is present in both our monomer and dimer structures. Our structural data suggest that disruption of the dimer should precede phosphorylation of the COR-B residues that lead to activation; the COR-B loop is relatively buried in the structure, surrounded by the ROC and LRR domains from the same monomer, and the ANK repeats that come from the other monomer to block the kinase (Figure 4H-K). Future studies will also determine whether, in addition to relieving autoinhibition, phosphorylation of the COR-B loop enables new interactions, with LRRK1 and/or other partners.

Although much remains to be done to understand how LRRK proteins are regulated, it is tempting to speculate that the apparently more stringent regulation of LRRK1 may be related to the fact that disease-linked mutations in LRRK1 are autosomal recessive loss-of-function, in contrast to those in PD-linked LRRK2. It may be that a hyperactive LRRK1 is more acutely deleterious than a hyperactive LRRK2 in cells.

### Structural signatures of LRRK1 and LRRK2

Understanding how LRRK1 and LRRK2 have similar domain architectures and cellular functions, yet very different disease associations and genetics is key to deciphering the etiology of the diseases with which they are linked. One of our goals in this study was to determine what structural features are unique to LRRK1 or shared between LRRK1 and LRRK2. LRRK1-specific features would shed light on LRRK1’s involvement in bone diseases, while those specific to LRRK2 would shed light on its involvement in PD. Our structures of LRRK1 revealed that the apparent similarity in the domain organization of these two proteins belies very different autoinhibitory mechanisms. Our evolutionary analysis showed that two of the structural elements involved in LRRK1’s autoinhibition—a long loop in the WD40 domain and a COR-B loop that directly inhibits the kinase—are conserved in LRRK1s but absent in LRRK2s, suggesting that autoinhibition is central to LRRK1’s function. Interestingly, LRRK3s, including those found in arthropods (*e*.*g*., *Drosophila*) and nematodes (*e*.*g*., *Caenorhabditis*) lack the long WD40 loop but share an extended COR-B loop with LRRK1. While it is unclear what role these might play in LRRK3 function in these species, it does suggest that studies of LRRK protein function from model invertebrates should be treated with caution when extrapolating to human LRRKs. Conversely, our analysis showed that the LRRK2-specific basic patches found in its ROC domain, which mediate LRRK2’s interaction with microtubules, are absent in LRRK1s but conserved in LRRK2s, although the full complement of three basic patches is only found in jawed vertebrates. Although the physiological relevance of microtubule binding by LRRK2 remains to be established, our data suggest that the functional role of the basic patches in LRRK2 is unique to this LRRK. Given that the formation of microtubule-associated filaments requires two dimerization interfaces in LRRK2 (COR-B:COR-B and WD40:WD40), we were interested in determining whether their presence was conserved in LRRK2s. However, we were unable to do this as the residues in the WD40:WD40 interface are poorly conserved, even among vertebrate LRRK2s.

The last structural feature we analyzed that distinguishes LRRK1 and LRRK2 is the length of their αC helices, found in the N-lobe of the kinase. Unlike the features discussed before, for which we have shown specific functions, we do not yet understand the mechanistic implications of the length of the αC helix, though our evolutionary analysis indicated that a longer helix is a signature of LRRK1s and LRRK3s. Our structures of LRRK1 suggested that the more extensive interaction between a longer αC helix and a hydrophobic pocket in the COR-B domain may rigidify LRRK1 relative to LRRK2 (Figure S2). Future studies will determine whether LRRK1 and LRRK2 have different dynamics despite their similar domain architecture, and whether those are related to their function and regulation. One possibility is that increased rigidity in LRRK1 facilitates the formation of its autoinhibited dimer.

## Acknowledgements

This research was funded in part by Aligning Science Across Parkinson’s (grant number ASAP-000519) (S.K., S.L.R.-P., A.E.L.) through the Michael J. Fox Foundation for Parkinson’s Research (MJFF) (S.K., S.L.R.-P., A.E.L.). The work was also funded by the Howard Hughes Medical Institute (where S.L.R-P. is an Investigator); the Michael J. Fox Foundation (grant number 18321 to A.E.L. and S.L.R.-P.); and the National Institutes of Health (R35GM133633 to M.D.D.). We would like to thank Dario Alessi for sharing unpublished data on the phosphorylation sites in LRRK1’s COR-B domain. We also thank the UC San Diego Cryo-EM Facility, the Nikon Imaging Center at UC San Diego, and the UC San Diego Physics Computing Facility for IT support. For the purpose of open access, the authors have applied a CC BY 4.0 public copyright license to all Author Accepted Manuscripts arising from this submission.

## Author contributions

JMR and YXL performed all the cryo-EM work. SM, JMR and YXL prepared samples for structural studies. AD and RA carried out all cell-based assays. EJF and MDD carried out the evolutionary analyses. SK, MDD, SRP and AEL supervised the work. JMR, SRP, EJF, MDD and AEL wrote the manuscript, and JMR, AD, RA, SM, MDD, SRP and AEL edited it.

## Data Availability

Cryo-EM maps of the LRRK1 monomer (LRRK1(Δ1-19)) (EMD-27813), the LRRK1 dimer (LRRK1(FL)) (EMD-27817), a symmetry expanded monomer from the dimer map (EMD-27818), and local refinements of the monomer structure—around KW (EMD-27816), RCKW (EMD-27814), and RCK (EMD-27815)—were all deposited in the EM Data Bank. Models of the LRRK1 monomer (PDB: 8E04), LRRK1 dimer (PDB: 8E05), and symmetry expanded monomer from the dimer structure (PDB: 8E06) were all deposited in the Protein Data Bank.

## Materials and Methods

### Cloning and mutagenesis

For baculovirus expression, the DNA coding for LRRK1 was codon optimized for *Spodoptera frugipera* (Sf9) cells and synthesized by Epoch Life Science. The DNA was cloned via Gibson assembly into the pKL baculoviral expression vector, with an N-terminal His6-Z-tag and a TEV protease cleavage site. LRRK1 variants were generated using Q5 site-directed mutagenesis (New England Biolabs, NEB). The pKL plasmid was used for the generation of recombinant Baculoviruses using the Bac-to-Bac expression system (Invitrogen).

For mammalian expression, pcDNA5-FRT-TO-LRRK1 from MRC-PPU was used. The various LRRK1 mutants were generated using QuikChange sitedirected mutagenesis (Agilent), or Q5 site-directed mutagenesis (NEB) following manufacturer’s instructions. Plasmid design was performed using SnapGene software (Insightful Science; snapgene.com), and all plasmids were sequence-verified. EGFP-Rab7 was obtained from Addgene (#12605).

### LRRK1 expression and purification

N-terminally tagged His6-Z-TEV-LRRK1(FL) was expressed in Sf9 insect cells. Insect cells were infected with baculovirus and grown at 27°C for 3 days. Cells were harvested and cell pellets were resuspended in lysis buffer (50 mM HEPES pH 7.4, 500 mM NaCl, 20 mM imidazole, 0.5 mM TCEP, 5% glycerol, 5 mM MgCl2, 20 μM GDP, 0.5 mM Pefabloc and cOmplete EDTAfree protease inhibitor cocktail (Roche). Cells were lysed using a Dounce homogenizer and clarified by centrifugation. The supernatant was incubated for 1 hr with Ni-NTA agarose beads (Qiagen) equilibrated in lysis buffer. Beads were applied to a gravity column where they were extensively washed with lysis buffer, followed by elution in lysis buffer containing 300 mM imidazole. The eluted protein was diluted to 250 mM NaCl and loaded onto a SP Sepharose column (Cytiva) equilibrated in buffer (50 mM HEPES pH 7.4, 250 mM NaCl, 0.5 mM TCEP, 5% glycerol, 5 mM MgCl2, 20 μM GDP, 0.5 mM Pefabloc). The protein was eluted using a 250 mM to 2.5 M NaCl gradient. Fractions containing LRRK1(FL) were pooled, diluted to

∼500 mM NaCl, and incubated with TEV protease overnight at 4°C. The protein was concentrated and put directly over a S200 size exclusion column (Cytiva) equilibrated in storage buffer (50 mM HEPES pH 7.4, 150 mM NaCl, 0.5 mM TCEP, 5% glycerol, 5 mM MgCl2 and 20 μM GDP). The protein was concentrated to ∼5-6 μM and flash frozen in liquid nitrogen for storage. N-terminally tagged His6-Z-TEV-LRRK120-2015 was purified using the same protocol.

### Cryo-electron microscopy

*Sample preparation:* For LRRK1(Δ1-19), grid samples contained 2 μM LRRK1(Δ1-19), 3 μM Rab7a, purified as previously described (Snead et al., 2022), 1 mM GTP and 1 mM AMPcPP. For LRRK1(FL), 5.2 μM LRRK1(FL) was spiked with a final concentration of 0.06% brij-35 directly before vitrification. For both samples, 4 μL of sample was applied to a freshly plasma cleaned Ultrafoil grid (Electron Microscopy Sciences, EMS). Samples were blotted for 4s using a blot force of 20 followed by vitrification using a Vitrobot Mark IV (FEI) set to 4°C and 100% humidity.

*Data collection:* Cryo-EM data was collected on a Talos Arctica (FEI) operated at 200 kV and equipped with a K2 Summit direct electron detector (Gatan). Leginon (Suloway et al., 2005) was used for automated data collection. For the LRRK1(Δ1-19) dataset, we collected 1,310 movies at a nominal magnification of 36,000x and object pixel size of 1.16 Å. Movies were dose-fractionated into 200 ms frames for a total exposure of 12s with a dose rate of ∼5.1 electrons Å-2s-1. For the LRRK1(FL) sample, multiple datasets were collected and combined as the addition of detergent resulted in very few particles per image. Movies were collected with the same parameters as the LRRK1(Δ1-19) dataset.

*LRRK1(Δ1-19) monomer reconstruction:* cryoSPARC Live (Punjani et al., 2017b) was used to align movie frames (patch motion correction) and estimate the CTF (patch CTF estimation). Micrographs with a CTF worse than 6 Å were removed. Particles were picked using crYOLO (Wagner et al., 2018) with a previously trained model on the dose-weighted images. Particles were extracted with a down-sampled pixel size of 4.64 Å/px and used in multiple rounds of 2D classification in Relion 3.0 (Zivanov et al., 2018, p. 3). Particles were extracted to 1.16 Å/px and all subsequent processing was done in cryoSPARC (Punjani et al., 2017b). Particles were subjected to ab initio reconstruction and the best two classes were combined and used in non-uniform refinement. To further separate out heterogeneity, another round of ab initio reconstruction was performed. Heterogeneous refinement was then done to separate particles with and without the LRR domain. 2D classification was carried out on each resulting subset to remove lingering dimer particles or recover good particles. Particles were then combined and put through a last round of non-uniform refinement for a final resolution of 3.7 Å.

*LRRK1(FL) dimer reconstruction:* cryoSPARC Live (Punjani et al., 2017b) was used to align movie frames (patch motion correction) and estimate the CTF (patch CTF estimation). Particles were picked using crYOLO (Wagner et al., 2018) trained with manual particle picks. Particles were extracted with a down-sampled pixel size of 4.11 Å/px and used in multiple rounds of 2D classification implemented in cryoSPARC (Punjani et al., 2017b). Good particles were subjected to ab initio reconstruction. The best two classes were combined, and the particles were used in heterogeneous refinement. Final particles were taken and used in non-uniform refinement with C2 symmetry and optimized per-group CTF params enabled for a final resolution of 4.6 Å. Symmetry expansion followed by local refinement resulted in a monomer structure of 3.6 Å.

*Model building for monomeric LRRK1(Δ1-19):* The AlphaFold (Jumper et al., 2021) model of human LRRK1, Q38SD2, was docked into the LRRK1(Δ1-19) map. This model has the kinase modeled in the closed conformation, and so we used Phenix (Afonine et al., 2018) real space refine to rigid body fit each domain into the map. Discrepancies in the AlphaFold model were manually corrected in COOT (Emsley and Cowtan, 2004). To aid in model building, local refinements with masks around the KW domains (EMDB 27816), RCKW domains (EMDB 27814) and RCK domains (EMDB 27815) were performed. These maps only minimally improved the local resolution (∼0.1-0.3 Å), and therefore specific models for these maps were not included. Model refinement was done using a combination of Phenix real space refine (Afonine et al., 2018) and Rosetta Relax (ver 3.13) (Wang et al., 2016).

*Model building for dimeric LRRK1(FL):* The monomer model for LRRK1RCKW and the ANK-LRR domains from AlphaFold (Jumper et al., 2021) model Q38SD2 were docked into the dimer symmetry expansion map and rigid body fit using Phenix (Afonine et al., 2018) real space refine. The model was iteratively built in COOT (Emsley and Cowtan, 2004) and refined using Phenix (Afonine et al., 2018) real space refine and Rosetta Relax (ver 3.13) (Wang et al., 2016).

### Cell Culture and transfection

HEK293T cells were obtained through ATCC (CRL-3216). Cells were cultured in DMEM growth medium (Gibco) supplemented with 10% fetal bovine serum (Gibco) and 1x penicillin/streptomycin solution (Gibco). Cells were plated on 6-well dishes (200,000 cells per well) 24 h before transfection. Each culture well was transfected with 1 μg of FLAG-LRRK1 construct and 500 ng of EGFP-Rab7 using polyethylenimine (Polysciences) and cultured at 37°C with 5% CO2 for 36 h before harvesting.

### Western blot analysis and antibodies

For western blot quantification of LRRK1 protein expression and Rab7 phosphorylation, cells were harvested by scraping, rinsed with ice-cold 1x PBS, pH 7.4 and lysed on ice in RIPA buffer (50 nM Tris pH7.5, 150 mM NaCl, 0.2% Triton X-100, 0.1% SDS, with cOmplete EDTA-free protease inhibitor cocktail (Sigma-Aldrich) and PhoStop phosphatase inhibitor (Sigma-Aldrich). Lysates were rotated for 15 min at 4°C and clarified by centrifugation at maximum speed in a 4°C microcentrifuge for 15 min. Supernatants were then boiled for 10 minutes in SDS buffer. Replicates were performed on independently transfected cultures.

Lysates were run on 4-12% polyacrylamide gels (NuPage, Invitrogen) for 50 minutes at 180V and transferred to polyvinylidene difluoride (Immobilon-FL, EMD Millipore) for 4 h at 200 mA constant current. Blots were rinsed briefly in MilliQ water and dried at room temperature for at least 30 min. Membranes were briefly reactivated with methanol and blocked for 1 h at room temperature in 5% milk (w/v) in TBS. Antibodies were diluted in 1% milk in TBS with 0.1% Tween-20 (TBST). Primary antibodies used for immunoblots were as follows: mouse anti-GFP (Santa Cruz, 1:2500 dilution), rabbit anti-LRRK1 (Abcam, 1:500 dilution), rabbit anti-GAPDH (Cell Signaling Technology, 1:3000 dilution), and rabbit anti-phospho-S72Rab7A (MJF-38, 1:1000 dilution). Secondary antibodies (1:10,000) used for western blots were IRDye 680RD Goat anti-Mouse and IRDye 780RD Goat anti-Rabbit (Li-COR). Primary antibodies were incubated overnight at 4°C, and secondary antibodies were incubated at room temperature for 1 h. For quantification, blots were imaged on an Odyssey CLx Imaging System (Li-COR) controlled by Imaging Studio software (v.5.2) (Li-COR), and intensity of bands quantified using Image Studio Lite software (v.5.2). All statistical analyses were performed in GraphPad Prism (9.3.1; GraphPad Software).

### Phylogenetic analyses

Human LRRK1 (accession NP_078928.3) and LRRK2 (accession NP_940980.4) were used as a BLASTp (Altschul et al., 1990) search query against the Reference Sequence (RefSeq) protein database with an e-value cutoff of 1e-20 and a query coverage cutoff of 40%. Resulting sequences were aligned using Clustal Omega (Sievers et al., 2011) and duplicate and poorly aligning sequences were removed. To reduce the number of nearly identical sequences, sequences with >80% sequence identity were reduced to a single unique sequence using CD-HIT (Fu et al., 2012) with a 0.8 sequence identity cutoff. The resulting 273 sequences (accession numbers and species names found in Supplemental Dataset 1) were realigned with Clustal Omega using two rounds of iteration to optimize the alignment throughout the sequences. The resulting alignment is found in Supplemental Dataset 2. To generate maximum likelihood phylogenetic trees of LRRK proteins, IQ-TREE (Nguyen et al., 2015) phylogenies were generated using the “-bb 1000 -alrt 1000” commands for generation of 1000 ultrafast bootstrap (Hoang et al., 2018) and SH-aLRT support values. The best substitution model (JTT+F+I+G4) was determined by ModelFinder (Kalyaanamoorthy et al., 2017) using the “-m AUTO” command. To confirm that the phylogenetic inferences were not influenced by regions of LRRK proteins that are not well conserved across all family members, the ROC-CORA region (corresponding to human LRRK1 residues 632-995) and kinase domain region (corresponding to human LRRK1 residues 1229-1534) were extracted from the alignment and concatenated and used as input for IQ-TREE. The resulting phylogeny, using the best substitution model (Q.insect+I+G4), shows similarly strong support values in major branches of the phylogeny (Figure S6). Complete phylogenetic trees, with support values, can be found in Supplemental Dataset 3.

To determine length of WD40 and COR-B loops, and the aC helix region, well-aligning regions of the alignment, which often corresponded to ordered regions of the LRRK1 and LRRK2 structures, were used as boundaries elements to count the number of intervening residues. Boundary amino acid sequences and residue numbers are shown in Figure 7D. For the COR-B loop in cnidarian LRRK4, the automated alignment did not identify the well conserved WxxGfxf C-terminal boundary element because it is 200+ residues farther from the N-terminal boundary element in cnidarian LRRK4s than in any other LRRK proteins. To measure the loop length, this well conserved WxxGfxf motif was manually identified in cnidarian LRRK4s and used to count the intervening COR-B “loop” region. To determine the presence of basic patches, well-aligning boundary elements flanking each LRRK2 basic patch were identified as described above. For each sequence shown in figure 7G, intervening sequences between the indicated boundary elements were manually searched for any occurrence of three basic residues within a four-residue window.

Consensus logos and alignment visualization for basic patch 3 and the COR-B loop region were generated using Geneious Prime 2022.1.1 (https://www.geneious.com).

---

**Supplementary Table 1.**
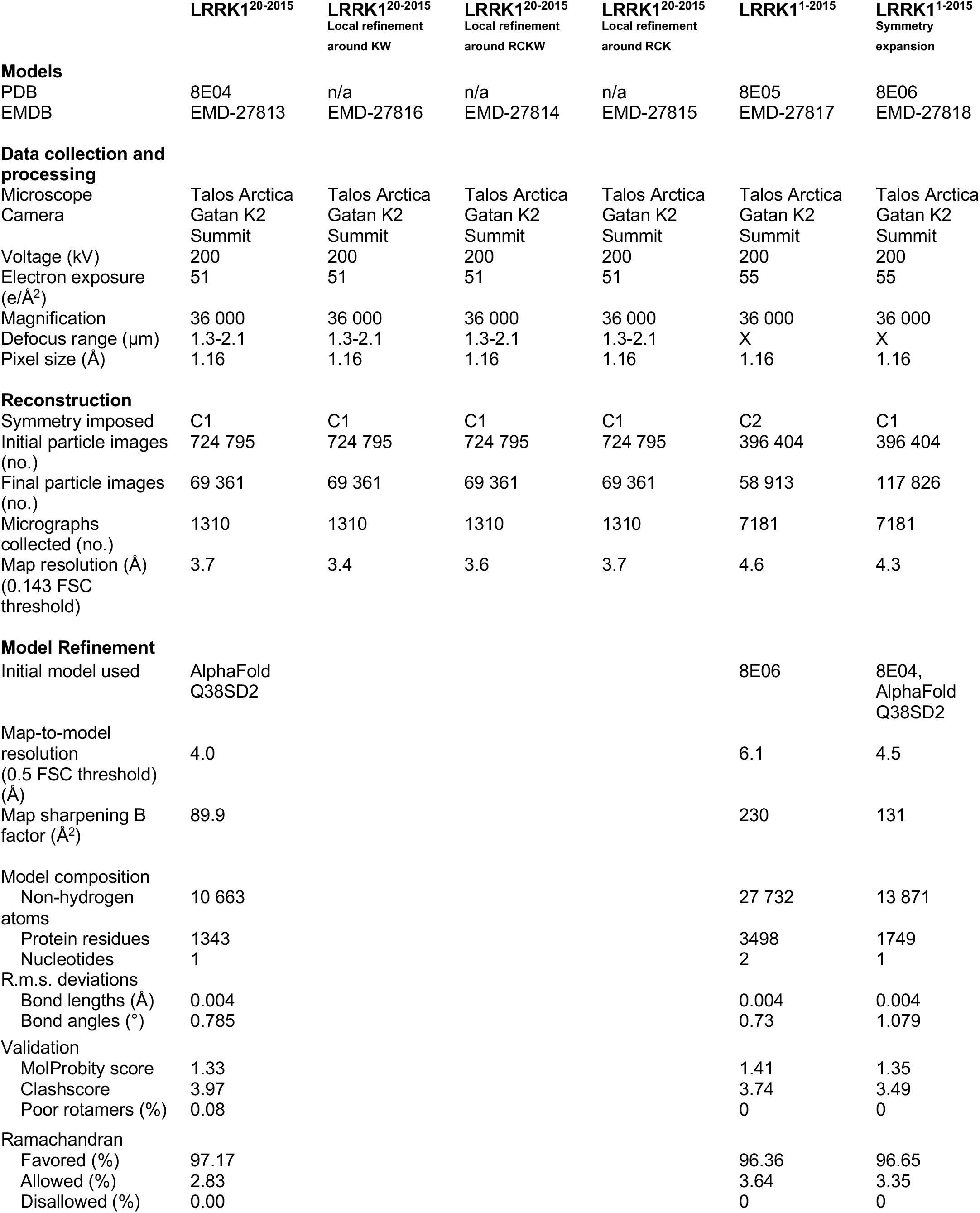
Cryo-EM data collection, refinement, and validation statistics. Related to Figures S1 and S4.

**Figure S1.**
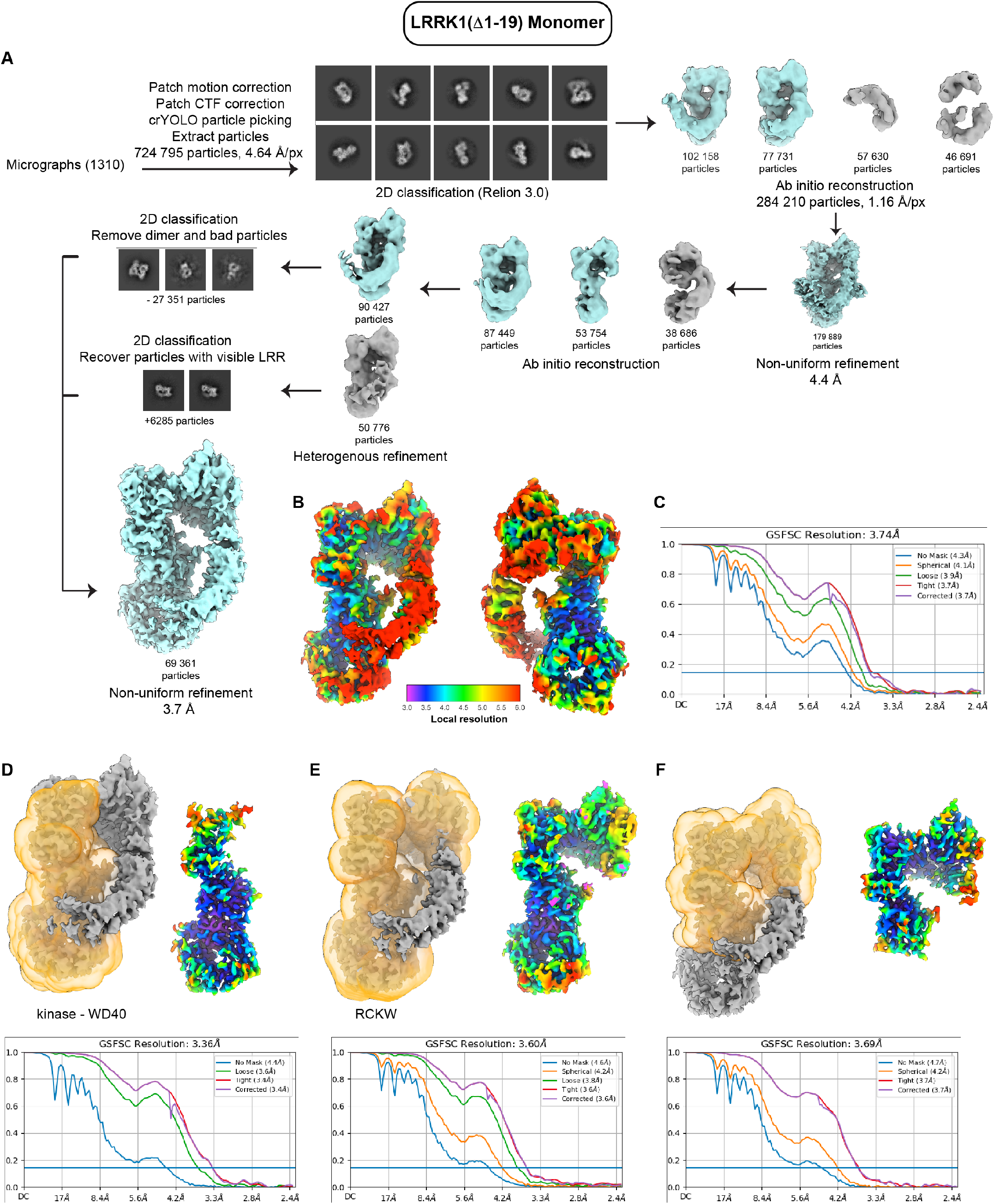
Cryo-EM workflow for monomeric LRRK1. **(A)** Data processing and reconstruction of monomeric LRRK1. **(B)** Map of monomeric LRRK1 colored by local resolution. **(C)** FSC plot for monomeric LRRK1. **(D-F)** Local refinements of (D) CORB-kinase-WD40, (E) ROC-CORB-kinase-WD40 and (F) ROC-CORB used in model building with corresponding FSC plots and maps colored by local resolution.

**Figure S2.**
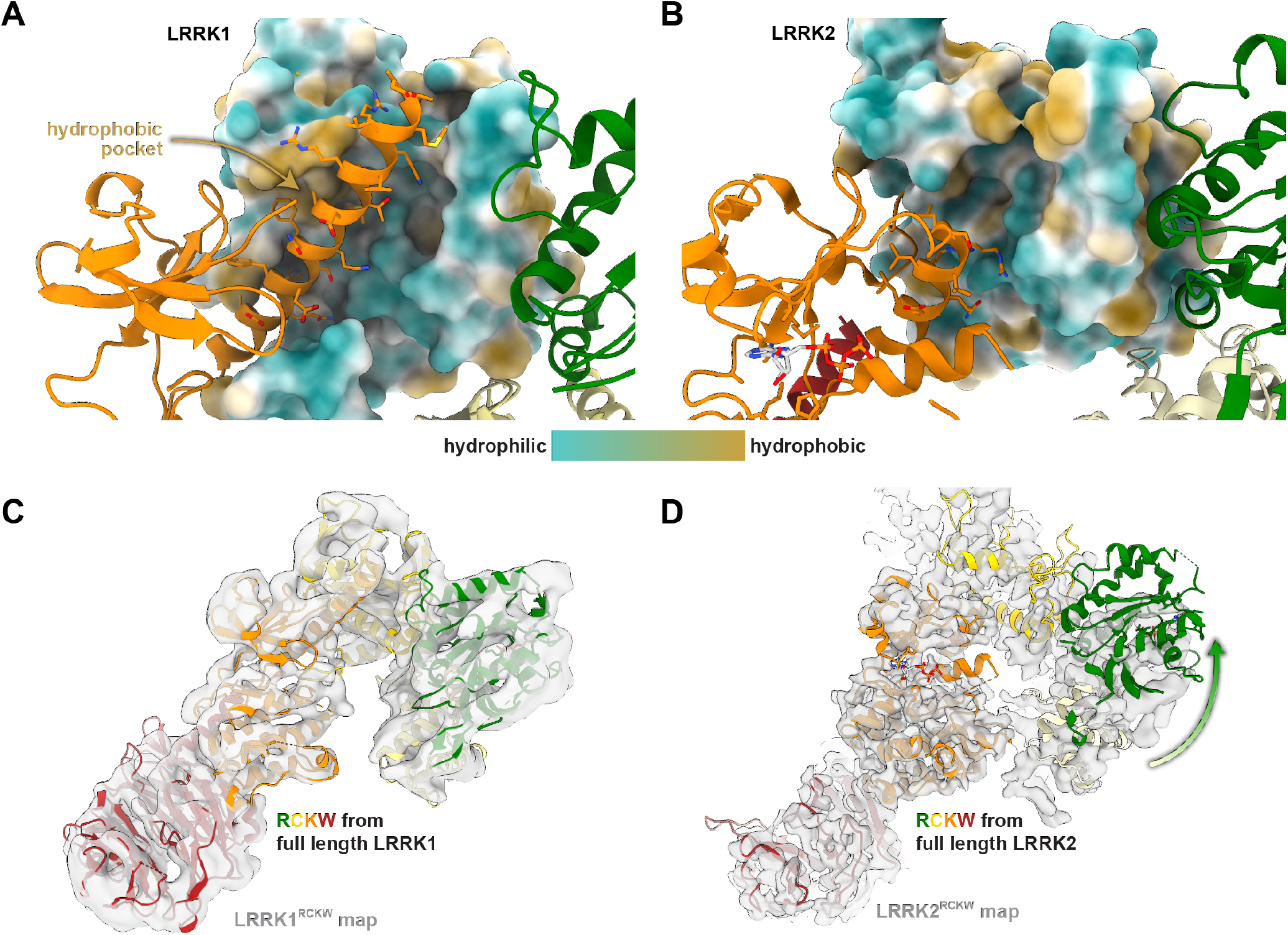
Interface between LRRK1’s C-alpha helix and the COR-B domain. **(A)** The N-lobe of LRRK1’s kinase domain is shown in ribbon representation, while the COR-B domain is shown in surface representation colored by its hydrophilicity/hydrophobicity. A hydrophobic pocket in the COR-B domain is indicated, and the side chains for residues in the C-alpha helix are shown. **(B)** Equivalent view for LRRK2. **(C**,**D)** The RCKW portion of full-length LRRK1 (C) or LRRK2 (D) were docked into cryo-EM maps of LRRK1RCKW (C) or LRRK2RCKW (D). The arrow in (D) indicates that the ROC domain from full-length LRRK2 is rotated relative to its position in the cryo-EM map of LRRK2RCKW.

**Figure S3.**
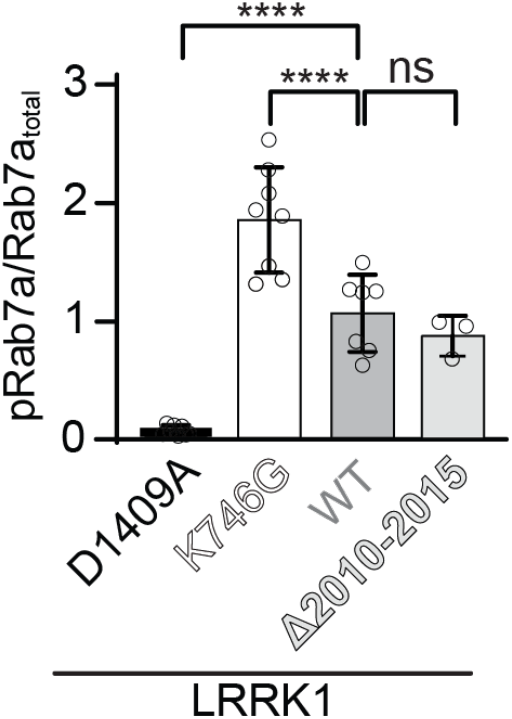
Deleting the last 6 residues of LRRK1 has no effect on Rab7a phosphorylation in cells. Rab7a phosphorylation in 293T cells expressing GFP-Rab7a and full-length WT LRRK1 or LRRK1(Δ2010-2015), where the last 6 residues were deleted. LRRK1(K746G), which is known to increase Rab7 phosphorylation in cells, and LRRK1(D1409A), which is known to be kinase inactive, were tested as well. 293T cells were transiently transfected with the indicated plasmids encoding FLAG-LRRK1 (wild type or mutant) and GFP-Rab7. Thirty-six hours post-transfection the cells were lysed, immunoblotted for phosphor-Rab7 (pS72), total GFP-Rab7, and total LRRK1. The mean ± s.e.m. is shown. ****p<0.0001, ns=not significant, one-way ANOVA Individual data points represent separate populations of cells obtained across at least three independent experiments (n ≥ 3).

**Figure S4.**
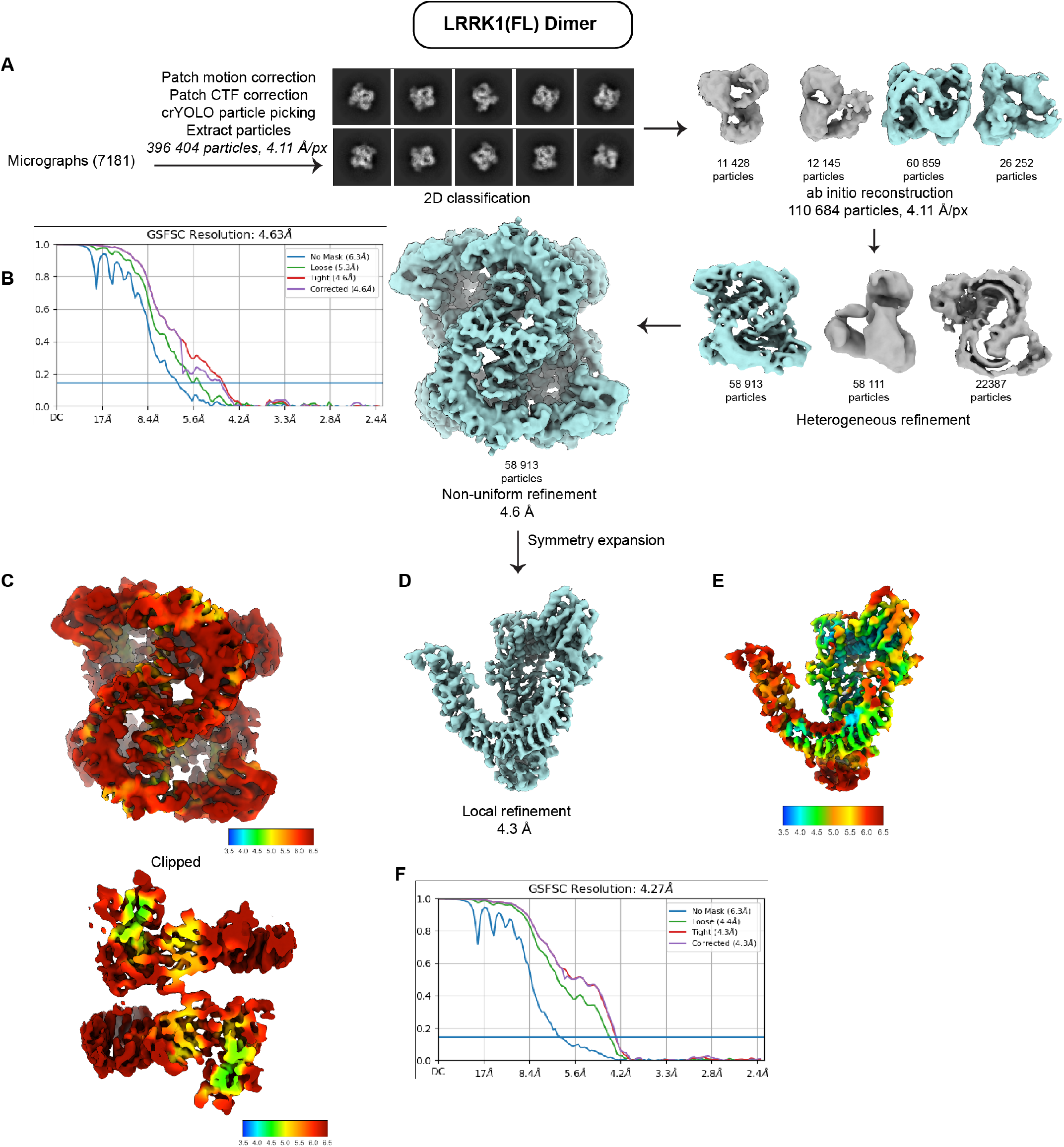
Cryo-EM workflow for dimeric LRRK1. **(A)** Data processing and reconstruction of dimeric LRRK1. **(B)** FSC plot for dimeric LRRK1. **(C)** Dimer map colored by local resolution (top), with clipped surface (bottom) showing higher resolution in the interior of the map. **(D)** Particles were symmetry expanded and used in local refinement to obtain a 4.3Å reconstruction of a LRRK1 monomer. **(E)** Symmetry expanded map colored by local resolution. **(F)** FSC plot for the symmetry expansion map.

**Figure S5.**
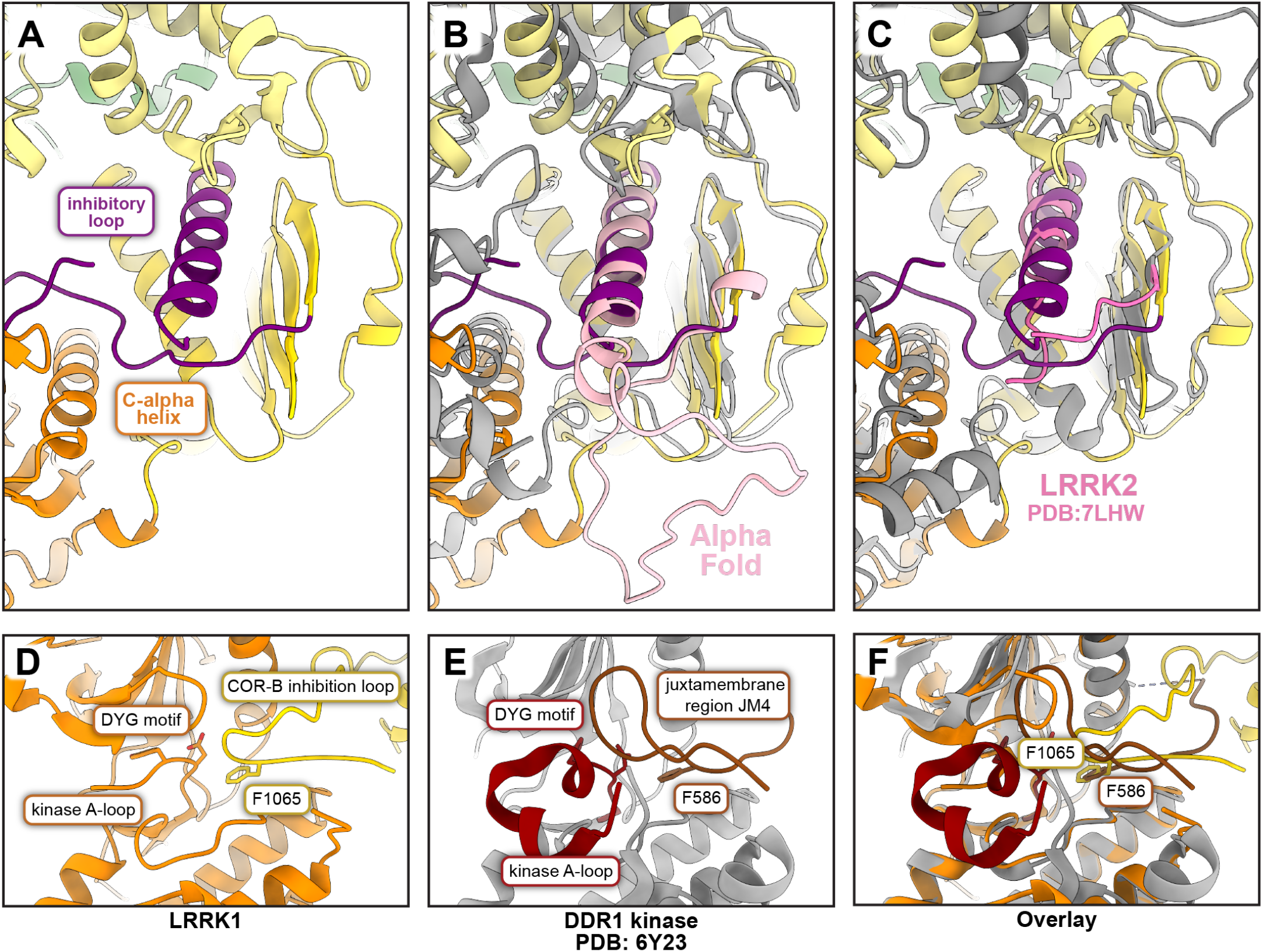
LRRK1’s COR-B autoinhibitory loop. **(A-D)** Close ups of the kinase-COR-B region of LRRK1. **(A)** The COR-B autoinhibitory loop is shown in dark purple. **(B)** Same as (A), with the AlphaFold model of LRRK1 shown in light purple. The portion corresponding to the autoinhibitory loop identified in our structure was modeled as disordered in the AlphaFold structure. **(C)** The model for full-length LRRK2 (PDB: 7LHW), in medium purple, is overlayed on the structure of LRRK1 shown in panel (A); the long COR-B loop present in LRRK1 is absent in LRRK2. **(D-F)** DDR1 kinase uses an autoinhibitory mechanism analogous to that of LRRK1. **(D)** View of the active site of LRRK1’s kinase, similar to Figure 6D. **(E)** Active site of DDR1 kinase (PDB: 6Y23). **(F)** Overlay of LRRK1 and DDR1. Note that F1065 in LRRK1 and F586 in DDR1 both occupy the same back pocket in the kinase’s active site.

**Figure S6.**
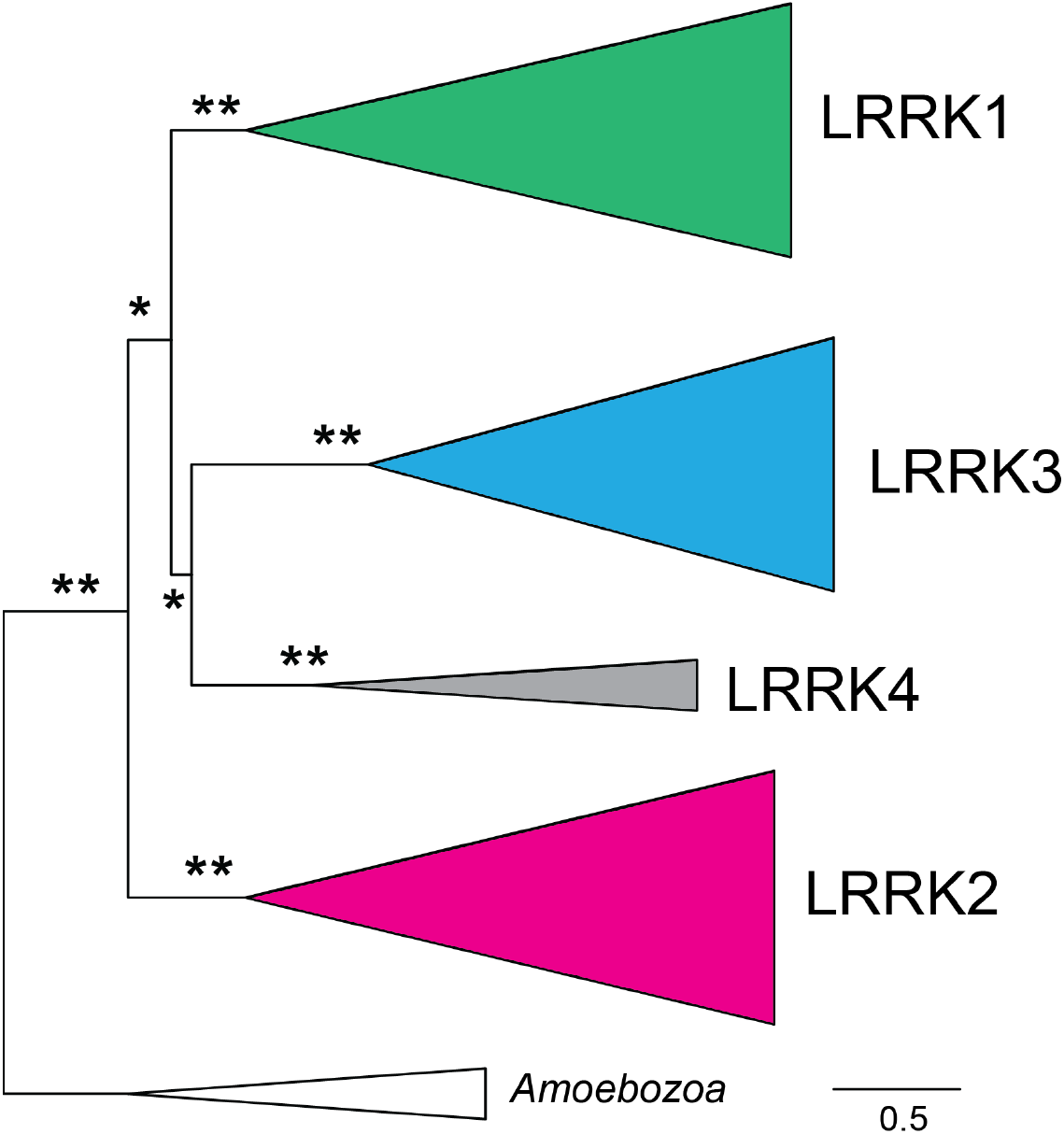
Maximum likelihood phylogenetic tree of LRRK protein homologs based only on ROC, COR-A, and kinase domain regions. An alignment of full length LRRK protein homologs (protein accessions and alignment in supplemental datasets 1 and 2, respectively) was generated and then alignment regions corresponding to the ROC-COR-A domains (human LRRK1 residues 632-995) and kinase domain region (human LRRK1 residues 1229-1534) were extracted and concatenated. This extracted alignment, which only contains regions that are shared across all LRRK protein homologs, was used as input to generate a maximum likelihood phylogenetic tree of LRRK protein homologs (complete tree in supplemental dataset 3). Shading and rooting are the same as in Figure 7. Asterisks indicate bootstrap branch support (* >75% support, ** 100% support). Support for major LRRK protein clades is similar to the phylogenetic tree shown in Figure 7, which was generated from the full-length protein alignment.

**Figure S7.**
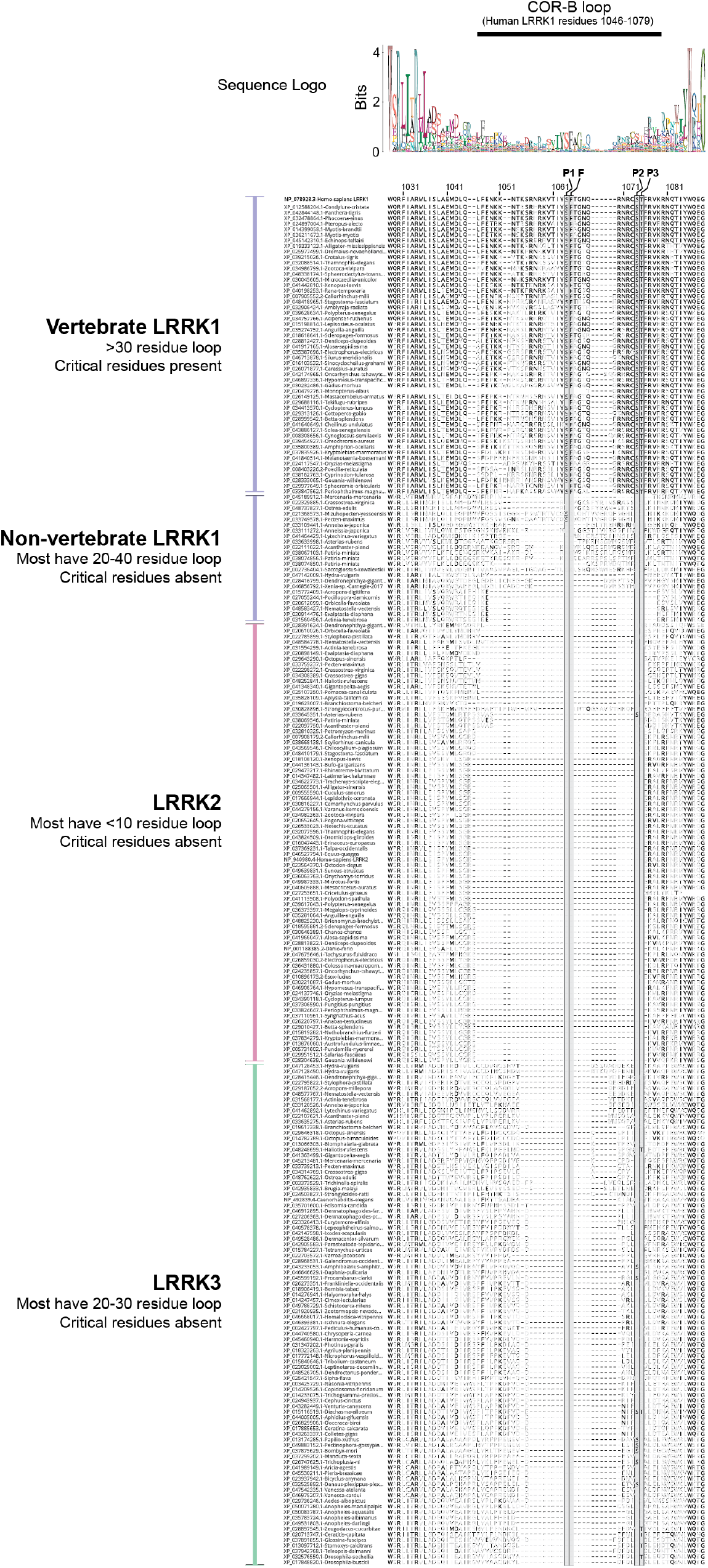
COR-B loop alignments in LRRK1, LRRK2, and LRRK3 proteins. Sequence alignment of the region of metazoan LRRK proteins that corresponds to the LRRK1 COR-B inhibitory loop (residues 1046-1079 in human LRRK1) and surrounding residues. The consensus sequence logo is shown at the top, with the alignment below. The Phe that occupies the back pocket of the kinase in the autoinhibited conformation (F1065 in human LRRK1) and the 3 sites of PKC phosphorylation (S1064, S1074, and T1075 in human LRRK1), are indicated above and highlighted in the alignment. Major species clades are shown at the left next to individual sequence accession numbers and species names. The human LRRK1 sequence is shown at the top of the alignment with residue numbers indicated. For other sequences in the alignment, residues in black are identical to human LRRK1. Residues in grey do not match the amino acid of human LRRK1, and dashes indicate there is no amino acid in this alignment position. The majority of vertebrate LRRK1 proteins have a COR-B loop >30 residues and retain the Phe and phosphorylation sites present in human LRRK1. Other LRRK proteins either have a “short” COR-B loop (*e*.*g*., LRRK2) or lack the Phe and phosphorylation sites.

**Figure S8.**
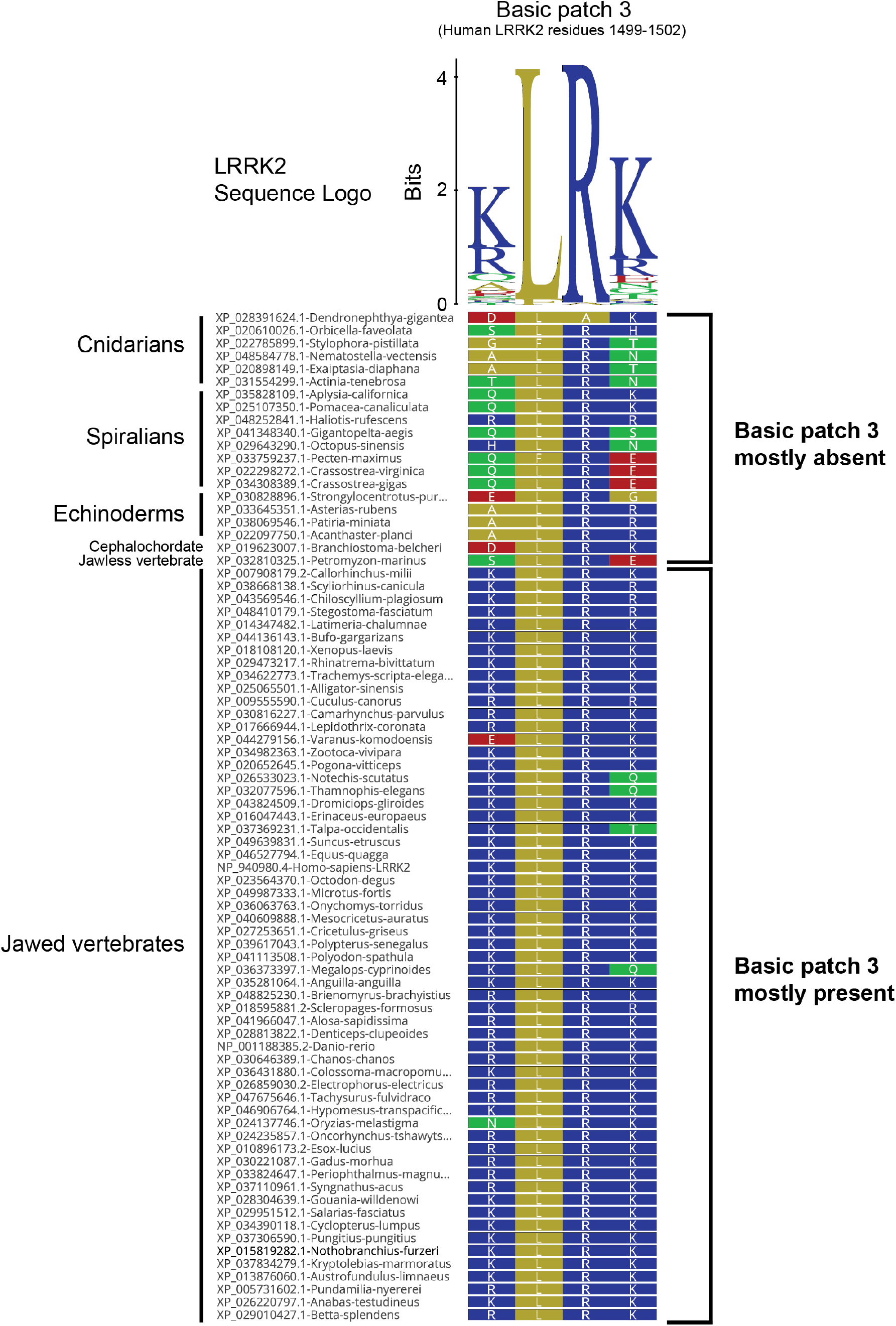
Presence of basic patch 3 in metazoan LRRK2s. Sequence alignment of the region of metazoan LRRK2 that corresponds to basic patch 3 (residues 1499-1502 in human LRRK2). The consensus sequence logo is shown at the top. Major species clades are shown at the left next to individual sequence accession numbers and species names. Amino acids are colored according to polarity, with basic residues shown in blue. Most jawed vertebrates have a basic-L-R-basic motif. Non-jawed vertebrate metazoans have a conserved central L-R motif, but the first and/or fourth residues tend to not be basic.

## SUPPLEMENTARY DATA

**Supplemental Dataset 1: Sequence accession numbers and species names for phylogenetic analyses**.

**Supplemental Dataset 2: LRRK protein alignment used for phylogenetic analyses**.

**Supplemental Dataset 3: Complete unrooted IQ-TREE phylogenetic trees for Figure 7 and Figure S6**.

For each node in the maximum likelihood tree, results from 1000 iteration SH-aLRT and 1000 iteration ultrafast bootstraps are shown as support values respectively.

