## Supplemental Dataset 3 for "Structure of LRRK1 and mechanisms of autoinhibition and activation"

### Supplemental File 3: Complete unrooted IQ-TREE phylogenetic trees for Figure 7 and Figure S6.

For each node in the maximum likelihood tree, results from 1000 iteration SH-aLRT and 1000 iteration ultrafast bootstraps are shown as support values respectively.

#### Full length LRRKs (tree shown in Figure 7).

Sequences: 273
Alignment length: 8025
Best fit model: JTT+F+I+G4

(((('XP_001134523.1-Dictyostelium-discoideum-AX4':0.2563764321000006,'XP_003284809.1-Dictyostelium-
purpureum':0.25727151260000003)'100/100':0.3053164368000001,('XP_012760012.1-Acytostelium-subglobosum-
LB1':0.5018290698999994,('XP_020428079.1-Heterostelium-album-PN500':0.33201488499999954,'XP_012749112.1-
Acytostelium-subglobosum-
LB1':0.5510420146000001)'94.5/100':0.10027851679999955)'100/100':0.2584922412999999)'100/100':1.5850425330000002,((('X
P_003294476.1-Dictyostelium-purpureum':0.2218172144999997,'XP_645923.1-Dictyostelium-discoideum-
AX4':0.22560306559999965)'100/100':0.36427902219999986,('XP_004361995.1-Cavenderia-
fasciculata':0.40284255510000033,('XP_020429574.1-Heterostelium-album-PN500':0.3135231597000008,'XP_012757658.1-
Acytostelium-subglobosum-
LB1':0.3591302831000007)'100/100':0.1662439184000002)'60.6/44':0.11059438669999988)'100/100':1.0702015119999997,('XP_
004352522.1-Acanthamoeba-castellanii-str.-Neff':1.4791149381999995,('XP_004353012.1-Acanthamoeba-castellanii-str.-
Neff':1.3845249063000002,(('XP_004337072.1-Acanthamoeba-castellanii-str.-Neff':0.3561119923000007,'XP_004339596.1-
Acanthamoeba-castellanii-str.-Neff':0.7332888954999994)'100/100':0.4159402818000002,('XP_004367915.1-Acanthamoeba-
castellanii-str.-Neff':0.7015831153000001,'XP_004367596.1-Acanthamoeba-castellanii-str.-
Neff':0.49956201440000036)'100/100':0.28340078260000023)'100/100':0.5388737897000002)'99.8/100':0.30591811010000036)'1
00/100':0.5072694200000001)'99.8/100':0.4442520272000001)'100/100':0.7249423630000003,((('XP_032810325.1-Petromyzon-
marinus':1.0797268855000004,(((('XP_041113508.1-Polyodon-spathula':0.21667165000000033,'XP_039617043.1-Polypterus-
senegalus':0.42142098770000036)'96.5/100':0.05817858220000005,(('XP_018595881.2-Scleropages-
formosus':0.3178158885000002,'XP_048825230.1-Brienomyrus-
brachyistius':0.3266565174)'100/100':0.11453074280000042,(((('XP_041966047.1-Alosa-
sapidissima':0.24549328490000022,'XP_028813822.1-Denticeps-
clupeoides':0.29814200610000086)'100/100':0.09501612230000056,('XP_030646389.1-Chanos-
chanos':0.5194928014000002,('NP_001188385.2-Danio-rerio':0.42090413110000036,(('XP_047675646.1-Tachysurus-
fulvidraco':0.28357523200000045,'XP_036431880.1-Colossoma-
macropomum':0.13199602060000082)'35.1/100':0.02174648549999958,'XP_026859030.2-Electrophorus-
electricus':0.24254178400000015)'100/100':0.10320102030000022)'99.5/100':0.06155672330000726)'99.7/100':0.061953775399
999245)'100/100':0.08028576170000079,((('XP_030221087.1-Gadus-morhua':0.49692257629999936,(('XP_033824647.1-
Periophthalmus-magnuspinnatus':0.43793645210000065,'XP_037110961.1-Syngnathus-
acus':0.32057344270000065)'92.6/98':0.044789614900000885,((('XP_028304639.1-Gouania-
willdenowi':0.27585383689999965,'XP_029951512.1-Salarias-
fasciatus':0.2675878897999997)'100/100':0.0534463051999996,((('XP_015819282.1-Nothobranchius-
furzeri':0.17859093220000055,('XP_037834279.1-Kryptolebias-marmoratus':0.10758350649999926,'XP_013876060.1-
Austrofundulus-
limnaeus':0.12720079710000043)'100/100':0.08322812930000012)'99.9/100':0.043152080400000514,'XP_024137746.1-Oryzias-
melastigma':0.27496052090000056)'100/100':0.04559916979999912,'XP_005731602.1-Pundamilia-
nyererei':0.18291250639999923)'32.8/95':0.014414622300000346)'99.4/100':0.02660570399999962,(('XP_029010427.1-Betta-
splendens':0.22229639829999925,'XP_026220797.1-Anabas-
testudineus':0.10215737390000079)'100/100':0.06914640059999932,('XP_034390118.1-Cyclopterus-
lumpus':0.09108127630000062,'XP_037306590.1-Pungitius-
pungitius':0.18844880279999998)'100/100':0.07245891620000044)'71.5/96':0.00792399710000069)'99.6/98':0.0343922970000001
3)'99.6/99':0.06434565320000019)'100/100':0.08736610449999915,'XP_046906764.1-Hypomesus-
transpacificus':0.2534497678999994)'94.7/100':0.03989265889999949,('XP_010896173.2-Esox-
lucius':0.17753973489999986,'XP_024235857.1-Oncorhynchus-
tshawytscha':0.09617970010000043)'100/100':0.0680483829000007)'100/100':0.12911062380000082)'100/100':0.1226427384999
9974,('XP_035281064.1-Anguilla-anguilla':0.19440156659999985,'XP_036373397.1-Megalops-
cyprinoides':0.24570189659999997)'100/100':0.08113399580000014)'87.8/100':0.03459192529999999)'100/100':0.390022882100
00025)'100/100':0.2765019735000003,((('XP_044136143.1-Bufo-gargarizans':0.26859780280000045,'XP_018108120.1-Xenopus-
laevis':0.24094652259999982)'100/100':0.26295061350000015,(((((('XP_037369231.1-Talpa-
occidentalis':0.5978207093000005,'XP_016047443.1-Erinaceus-
europaeus':0.19171846809999948)'100/100':0.07714006980000043,'XP_049639831.1-Suncus-
etruscus':0.1893370504000007)'99.9/100':0.05094623709999979,((((('XP_027253651.1-Cricetulus-
griseus':0.05887739009999926,'XP_040609888.1-Mesocricetus-
auratus':0.03149180499999993)'100/100':0.01928507589999917,('XP_049987333.1-Microtus-
fortis':0.04846244610000028,'XP_036063763.1-Onychomys-

torridus':0.049778371100000385)'9.5/98':0.003142815100003623)'100/100':0.1049748774000001,'XP_023564370.1-Octodon-
degus':0.1424823931999999)'98.6/100':0.017525519900000397,'NP_940980.4-Homo-sapiens-
LRRK2':0.05716990909999975)'99.8/100':0.021937503200000208,'XP_046527794.1-Equus-
quagga':0.04810323380000034)'71.2/99':0.01985149570000022)'100/100':0.13605280069999992,'XP_043824509.1-Dromiciops-
gliroides':0.19065292919999965)'100/100':0.14898755050000023,(((('XP_032077596.1-Thamnophis-
elegans':0.0725354238999978,'XP_026533023.1-Notechis-
scutatus':0.07847749219999933)'100/100':0.28922802580000084,('XP_020652645.1-Pogona-
vitticeps':0.10307221490000007,'XP_044279156.1-Varanus-
komodoensis':0.1613780010000001)'96.3/100':0.0209335531999999)'59.6/100':0.0149121213000008,'XP_034982363.1-Zootoca-
vivipara':0.18981143770000042)'100/100':0.11085519339999994,('XP_034622773.1-Trachemys-scripta-
elegans':0.11037456209999963,('XP_025065501.1-Alligator-sinensis':0.11888617159999981,('XP_009555590.1-Cuculus-
canorus':0.07575772499999989,('XP_030816227.1-Camarhynchus-parvulus':0.03532522999999976,'XP_017666944.1-
Lepidothrix-
coronata':0.024496362199999844)'98/100':0.019456514399999847)'100/100':0.13623907619999986)'99.7/100':0.0327227820999
9957)'40.9/100':0.019456974599999732)'100/100':0.06380794459999972)'100/100':0.1118810086000016,'XP_029473217.1-
Rhinatrema-bivittatum':0.3413583612000002)'4.6/56':0.037653258599999795)'100/100':0.12316839779999977,'XP_014347482.1-
Latimeria-chalumnae':0.2821482232000001)'99.9/100':0.0776268411999985)'100/100':0.1385632915999997,('XP_007908179.2-
Callorhinchus-milii':0.22746868369999973,('XP_038668138.1-Scyliorhinus-canicula':0.16422707779999968,('XP_048410179.1-
Stegostoma-fasciatum':0.0878707018,'XP_043569546.1-Chiloscyllium-
plagiosum':0.08163651450000042)'100/100':0.08433315060000002)'100/100':0.19058899419999964)'100/100':0.19438814100000
013)'100/100':0.6604745432000003)'100/100':0.8429958120999999,((('XP_029643290.1-Octopus-
sinensis':1.8025870933000006,((('XP_041348340.1-Gigantopelta-aegis':0.7600764265000004,'XP_048252841.1-Haliotis-
rufescens':0.5059966606000001)'99.1/100':0.1366617554999996,('XP_035828109.1-Aplysia-
californica':0.5136257880999997,'XP_025107350.1-Pomacea-
canaliculata':0.6074339623)'100/100':0.25659587689999963)'100/100':0.2551880772999997,('XP_033759237.1-Pecten-
maximus':0.7801563366000002,('XP_034308389.1-Crassostrea-gigas':0.20319049530000033,'XP_022298272.1-Crassostrea-
virginica':0.2102088177999999)'100/100':0.8432656685)'100/100':0.2672917888999997)'92.6/99':0.1505568110000004)'100/100':0
.4563212414000004,('XP_019623007.1-Branchiostoma-belcheri':0.832235045,('XP_030828896.1-Strongylocentrotus-
purpuratus':0.7042040724999996,('XP_033645351.1-Asterias-rubens':0.18468931469999994,('XP_038069546.1-Patiria-
miniata':0.09157791720000041,'XP_022097750.1-Acanthaster-
planci':0.09793124479999982)'100/100':0.10113052210000006)'100/100':0.4940662480000002)'100/100':0.4602409559999998)'9
9.3/99':0.14585176960000013)'5.5/73':0.09193891039999968,('XP_028391624.1-Dendronephthya-
gigantea':1.6957290687999995,(('XP_022785899.1-Stylophora-pistillata':0.3180880022999997,'XP_020610026.1-Orbicella-
faveolata':0.1977283545999997)'100/100':0.5684936356000003,('XP_048584778.1-Nematostella-
vectensis':0.5820730676999997,('XP_031554299.1-Actinia-tenebrosa':0.4632789520999996,'XP_020898149.1-Exaiptasia-
diaphana':0.6852864271000003)'100/100':0.2769742344999999)'100/100':0.21880346689999985)'100/100':0.3194176501000001)'
100/100':0.3872382662999998)'98.4/100':0.30642103300000034)'100/100':0.9534872343999998,((('XP_012555367.2-Hydra-
vulgaris':1.2011152458999996,('XP_047143158.1-Hydra-vulgaris':0.5844803585999996,('XP_047143281.1-Hydra-
vulgaris':0.32515932599999964,(('XP_047144213.1-Hydra-vulgaris':0.2698337500000001,'XP_047144101.1-Hydra-
vulgaris':0.2614185613999993)'97.5/100':0.07012254290000008,'XP_047143514.1-Hydra-
vulgaris':0.20412236670000006)'99.8/100':0.10657749649999992)'100/100':0.2516401602)'100/100':0.4698544299999998)'100/10
0':1.3164795580000002,('XP_031569514.1-Actinia-tenebrosa':2.5127410551999994,(('XP_028409574.1-Dendronephthya-
gigantea':0.4709194152,'XP_046847823.1-Xenia-sp.-Carnegie-
2017':0.6075515023999998)'100/100':0.8394140370000001,('XP_032226651.2-Nematostella-
vectensis':0.5811856181000001,('XP_031571669.1-Actinia-tenebrosa':0.4039646725999999,'XP_020891986.1-Exaiptasia-
diaphana':0.44683922749999994)'100/100':0.22936743609999954)'100/100':0.8392149879000002)'93.8/100':0.250305792299999
83)'27.5/91':0.19830749559999994)'100/100':1.179776136,(((((((('XP_035701600.1-Folsomia-
candida':1.1650942663999997,('XP_040578378.1-Lepeophtheirus-salmonis':0.7973347906999999,'XP_023326413.1-Eurytemora-
affinis':0.5082050918999999)'100/100':0.23138938980000034)'88.3/58':0.09999451829999995,((((((((('XP_013174285.1-Papilio-
xuthus':0.2660136351000002,(('XP_045530211.1-Pieris-brassicae':0.2822821038000001,'XP_041989149.1-Aricia-
agestis':0.2550810829000003)'16.3/91':0.03121600519999923,(('XP_032525892.1-Danaus-plexippus-
plexippus':0.2725591747999996,('XP_047542335.1-Vanessa-atalanta':0.01737463159999919,'XP_046975207.1-Vanessa-
cardui':0.025124976400000776)'100/100':0.11495014199999964)'7.4/83':0.03260081270000015,'XP_023937942.1-Bicyclus-
anynana':0.12704732059999913)'100/100':0.06959410870000049)'100/100':0.07774378549999916)'100/100':0.095049978700000
54,(('XP_037875629.1-Bombyx-mori':0.31244381239999974,'XP_037299202.1-Manduca-
sexta':0.1933575653999995)'99.8/100':0.06616720319999914,'XP_026747625.1-Trichoplusia-
ni':0.22535318329999932)'97.7/79':0.05100953590000046)'68/76':0.06468685459999968,'XP_049883152.1-Pectinophora-
gossypiella':0.25725710360000065)'100/100':0.9104653429999994,(((('XP_049531803.1-Anopheles-
darlingi':0.06955866319999959,('XP_050083787.1-Anopheles-aquasalis':0.02961702540000033,'XP_035783724.1-Anopheles-
albimanus':0.028031424799999982)'99/100':0.03253954849999996)'100/100':0.20680420019999968,'XP_050071280.1-
Anopheles-maculipalpis':0.12829273240000028)'99.8/100':0.09348625610000028,'XP_029736246.1-Aedes-
albopictus':0.1449356182999999)'100/100':0.2112460425,(('XP_017848820.1-Drosophila-
busckii':0.14369500409999958,'XP_032576550.1-Drosophila-
sechellia':0.11859546170000002)'100/100':0.17335818260000035,(('XP_037942768.1-Teleopsis-
dalmanni':0.17672012749999944,('XP_028897545.1-Zeugodacus-cucurbitae':0.17246020299999998,'XP_020713747.1-Ceratitis-
capitata':0.16465295110000078)'100/100':0.1403328086000002)'33.3/99':0.024951067700000884,('XP_037891855.1-Glossina-
fuscipes':0.15767151939999913,'XP_013097712.1-Stomoxys-
calcitrans':0.09912452060000021)'100/100':0.06345352669999915)'99.6/100':0.06884163229999984)'100/100':0.49410483459999
99)'100/100':0.2748633025)'100/100':0.18870240329999977,((('XP_014209526.1-Copidosoma-
floridanum':0.22500048890000013,'XP_014233075.1-Trichogramma-

pretiosum':0.22235147760000018)'54/97':0.050068878900000335,'XP_003425729.1-Nasonia-
vitripennis':0.06209835940000019)'100/100':0.1052461719000018,(((('XP_044005005.1-Aphidius-
gifuensis':0.27580467760000005,'XP_015116519.1-Diachasma-
alloeum':0.15186333990000023)'100/100':0.07705799849999995,'XP_043282449.1-Venturia-
canescens':0.10664018599999991)'99.8/99':0.03715163569999991,'XP_024943937.1-Cephus-
cinctus':0.09078345200000015)'26.9/93':0.01379080119999987,(('XP_017885653.1-Ceratina-
calcarata':0.3524641675,'XP_043263337.1-Colletes-
gigas':0.10614457949999956)'98.7/99':0.0373262931999958,'XP_026829906.1-Ooceraea-
biroi':0.19329470959999995)'97/99':0.03503626909999906)'99.6/97':0.05264274049999962)'100/100':0.4678034248999996)'87.9
/98':0.06016355179999966,('XP_044740580.1-Chrysoperla-carnea':0.2508663870000003,('XP_045460940.1-Harmonia-
axyridis':0.34693196260000025,((('XP_018323263.1-Agrilus-planipennis':0.2374751124000003,'XP_031347202.1-Photinus-
pyralis':0.18563853920000017)'99.2/100':0.04713352509999574,'XP_017772148.1-Nicrophorus-
vespilloides':0.17630771199999984)'25.9/81':0.027843003600000138,(('XP_048526705.1-Dendroctonus-
ponderosae':0.2460501284000003,'XP_023029002.1-Leptinotarsa-
decemlineata':0.14836779979999992)'99.7/100':0.0538353135999996,'XP_015840646.1-Tribolium-
castaneum':0.1200190928999979)'97.6/100':0.04304652470000255)'94.9/82':0.0602742524000017)'100/100':0.140982947200
00037)'100/100':0.11426255279999964)'99.8/99':0.07753444360000028,(((('XP_025421547.1-Sipha-
flava':0.6558318074000002,'XP_018906419.1-Bemisia-
tabaci':0.29757970669999967)'86/73':0.07558117030000044,(('XP_014276941.1-Halyomorpha-
halys':0.18932811009999995,'XP_014247457.1-Cimex-
lectularius':0.17224449259999997)'100/100':0.3789181127000001,'XP_046668017.1-Homalodisca-
vitripennis':0.23048396009999994)'78.8/73':0.04221940199999974)'100/100':0.08931962840000018,'XP_026273351.1-
Frankliniella-occidentalis':0.4059901386)'24.4/92':0.03866588350000022,('XP_046393381.1-Ischnura-
elegans':0.5085383507000003,('XP_021920935.1-Zootermopsis-nevadensis':0.18551046740000032,'XP_049788729.1-
Schistocerca-
nitens':0.19286123310000036)'100/100':0.10065251530000019)'85/95':0.04387753820000029)'99.9/96':0.07106006470000015)'98
.2/94':0.0833070455999998,'XP_002427797.1-Pediculus-humanus-
corporis':0.6875348997000001)'100/100':0.18098012070000014,(('XP_046646629.1-Daphnia-
pulicaria':0.6306388651999999,'XP_043233053.1-Amphibalanus-
amphitrite':0.6289594653000004)'41.7/92':0.08212750499999988,'XP_045599192.1-Procambarus-
clarkii':0.48975863119999996)'47.4/54':0.06087395930000028)'76/50':0.045753778899999986)'100/99':0.14798906469999995,(((('
XP_027206363.1-Dermatophagoides-pteronyssinus':0.08616326290000043,'XP_046912895.1-Dermatophagoides-
farinae':0.07639294670000041)'100/100':1.0610387719999999,'XP_015784227.1-Tetranychus-
urticae':0.6201377422999999)'100/100':0.23018923520000012,(('XP_028968531.1-Galendromus-
occidentalis':0.1699824941000001,'XP_022703572.1-Varroa-
jacobsoni':0.17326415269999984)'100/100':0.8027386844000004,('XP_042147598.1-Ixodes-
scapularis':0.2724446925999997,'XP_049528486.1-Dermacentor-
silvarum':0.5682226022999997)'100/100':0.17995128609999966)'100/100':0.22994727699999995)'17.4/53':0.07614341010000025
,'XP_042905583.1-Parasteatoda-
tepidariorum':0.45351053080000003)'95.5/77':0.11475544069999977)'100/100':0.25857980909999956,('XP_003373529.1-
Trichinella-spiralis':0.9720460645999998,('XP_042935833.1-Brugia-malayi':0.6676544704000005,('NP_492839.4-Caenorhabditis-
elegans':1.1532923674999997,'XP_024503827.1-Strongyloides-
ratti':1.4189621424000007)'99.6/100':0.24008998419999994)'100/100':0.3964658433999997)'100/100':0.3842375413000001)'100/
100':0.29622491449999977,(('XP_029646318.1-Octopus-sinensis':0.003101162500000143,'XP_014782789.1-Octopus-
bimaculoides':0.021294832799999774)'100/100':1.3908911660000003,(('XP_045213481.1-Mercenaria-
mercenaria':1.1950190454999996,('XP_033739213.1-Pecten-maximus':0.7964298694999998,('XP_048762622.1-Ostrea-
edulis':0.17387903230000035,'XP_034314769.1-Crassostrea-
gigas':0.1627809065000001)'100/100':0.5667623129999999)'100/100':0.38877472430000015)'83/100':0.13870691710000038,('XP
_013066303.1-Biomphalaria-glabrata':0.8685263945999999,('XP_041363499.1-Gigantopelta-
aegis':0.7123373707000002,'XP_048248699.1-Haliotis-
rufescens':0.5864317762000004)'100/100':0.21253719689999961)'100/100':0.3368393946000001)'99.3/100':0.1749487403999996
4)'100/100':0.6026136506000004)'99.9/100':0.20658436140000003,('XP_019617338.1-Branchiostoma-
belcheri':1.2643110815999998,('XP_033120526.1-Anneissia-japonica':0.6484239218000001,('XP_041462892.1-Lytechinus-
variegatus':0.5262107758000001,('XP_022107621.1-Acanthaster-planci':0.1905019808999997,'XP_033635275.1-Asterias-
rubens':0.24991337400000013)'100/100':0.39733293030000016)'100/100':0.30584303839999993)'100/100':0.6682420374999998)'
97.6/100':0.1632168532999998)'99.9/100':0.254517452,(('XP_047128450.1-Hydra-
vulgaris':0.30354399500000007,'XP_047128453.1-Hydra-
vulgaris':0.24006485009999956)'100/100':1.3103168080000005,('XP_028415446.1-Dendronephthya-
gigantea':0.9185457829999999,(('XP_022795822.1-Stylophora-pistillata':0.2206527587,'XP_029187052.2-Acropora-
millepora':0.2174493552000003)'100/100':0.2736312854999996,('XP_031568177.1-Actinia-
tenebrosa':0.2396078080999997,'XP_048577767.1-Nematostella-
vectensis':0.23875462029999994)'100/100':0.20067519749999985)'100/100':0.33007123139999983)'100/100':0.30063199549999
986)'100/100':0.43688004589999974)'100/100':1.0698292,(('XP_047142009.1-Hydra-
vulgaris':1.5594693514000006,(('XP_046856792.1-Xenia-sp.-Carnegie-2017':0.39310886330000017,'XP_028416799.1-
Dendronephthya-gigantea':0.3483283502000001)'100/100':0.8667932336000002,(('XP_015772409.1-Acropora-
digitifera':0.34664500149999977,('XP_020612099.1-Orbicella-faveolata':0.17276194809999978,'XP_027055244.1-Pocillopora-
damicornis':0.29327961859999974)'94.1/100':0.09137317420000013)'100/100':0.30721590080000016,('XP_048583427.1-
Nematostella-vectensis':0.4853995257000001,('XP_020914476.1-Exaiptasia-diaphana':0.3865275643999997,'XP_031560456.1-
Actinia-
tenebrosa':0.26884383440000015)'100/100':0.17024721519999986)'100/100':0.23833447280000009)'100/100':0.42354973349999

98)'99.2/100':0.21579165470000028)'100/100':0.2923151070999994,(('XP_002736404.1-Saccoglossus-
kowalevskii':1.2733474563999998,(('XP_033109441.1-Anneissia-japonica':1.8863674867000002,('XP_041464429.1-Lytechinus-
variegatus':1.6125160937,('XP_033633998.1-Asterias-rubens':0.5477026083999998,((('XP_038074856.1-Patiria-
miniata':0.18500752889999994,'XP_038074850.1-Patiria-
miniata':0.2042754975000003)'100/100':0.46093926570000043,'XP_022111022.1-Acanthaster-
planci':0.2449506329000002)'39.8/99':0.05577470879999957,'XP_038067103.1-Patiria-
miniata':0.3425766961000001)'100/100':0.23013809259999984)'100/100':0.6843792586999999)'100/100':0.3667126534999996)'1
7.1/92':0.11621989919999987,'XP_033111272.1-Anneissia-
japonica':1.6207447599)'100/100':0.4585659074999997)'74.5/51':0.1503362667000001,(('XP_045189912.1-Mercenaria-
mercenaria':1.3855906107999996,(('XP_022329885.1-Crassostrea-virginica':0.20199702970000022,'XP_048737827.1-Ostrea-
edulis':0.16325060910000033)'100/100':0.8732151043999998,('XP_021368573.1-Mizuhopecten-
yessoensis':0.23693145910000002,'XP_033749578.1-Pecten-
maximus':0.1962011645999997)'100/100':0.9098066786999999)'99.9/100':0.3598977725000001)'100/100':0.8058546654000001,((
((('XP_040198253.1-Rana-temporaria':0.18458939099999938,'XP_041442810.1-Xenopus-
laevis':0.21067248760000012)'100/100':0.2725488615999998,'XP_030045605.1-Microcaecilia-
unicolor':0.23522048159999986)'96.8/100':0.05990050410000695,(((((('XP_024897004.1-Pteropus-
alecto':0.18103209769999928,('XP_014399058.1-Myotis-brandtii':0.06892369189999936,'XP_036211673.1-Myotis-
myotis':0.013808394099999788)'100/100':0.08280049529999989)'95.9/100':0.022159996300000095,'XP_012588204.1-Condylura-
cristata':0.09131006049999968)'42.7/51':0.009578981099999773,('XP_042844148.1-Panthera-
tigris':0.04395913940000007,'XP_032478864.1-Phocoena-
sinus':0.0690177043000002)'9.7/63':0.007225735799999633)'75.3/77':0.019536141000000562,'NP_078928.3-Homo-sapiens-
LRRK1':0.06794057060000025)'84.4/96':0.025614170499999922,'XP_045142310.1-Echinops-
telfairi':0.1674749345000004)'100/100':0.19507057329999977,(((('XP_032088514.1-Thamnophis-
elegans':0.07285537189999935,'XP_039215026.1-Crotalus-
tigris':0.06007232940000051)'100/100':0.1978605443000001,'XP_034986799.1-Zootoca-
vivipara':0.14983070200000004)'95.3/100':0.0356835537000002,'XP_048338174.1-Sphaerodactylus-
townsendi':0.2056701279000004)'100/100':0.1134008491999996,('XP_019333123.1-Alligator-
mississippiensis':0.11200223900000061,'XP_025977499.1-Dromaius-
novaehollandiae':0.10526335130000053)'99.9/100':0.04552891379999924)'93.7/100':0.04377777010000017)'99.4/100':0.0585281
57800000535)'100/100':0.16831260729999986,('XP_039628634.1-Polypterus-
senegalus':0.35712954960000065,(((((((('XP_016103532.1-Sinocyclocheilus-grahami':0.06495441820000014,'XP_026071877.1-
Carassius-auratus':0.06472534210000003)'100/100':0.1695840715000001,('XP_046713878.1-Silurus-
meridionalis':0.1951452966999998,'XP_035387696.1-Electrophorus-
electricus':0.1890264730000002)'98.2/100':0.04979906440000015)'100/100':0.10151518920000058,('XP_028812427.1-Denticeps-
clupeoides':0.23974209030000004,'XP_041917165.1-Alosa-
sapidissima':0.1799908328000006)'97.9/100':0.04948880300000204)'99.8/100':0.05994508579999991,(((((((('XP_033847562.1-
Periophthalmus-magnuspinnatus':0.6194979601999995,'XP_029977649.1-Sphaeramia-
orbicularis':0.17468427249999952)'98.7/80':0.061387804799999834,'XP_028333065.1-Gouania-
willdenowi':0.31247122039999997)'75.1/72':0.023548378300000117,(('XP_008308656.1-Cynoglossus-
semilaevis':0.24249856069999964,'XP_043886127.1-Solea-
senegalensis':0.20651294249999985)'100/100':0.08779628929999994,'XP_041640649.1-Cheilinus-
undulatus':0.2171602638000003)'96.4/80':0.020917408899999934)'94.4/79':0.017862989300000187,((((('XP_024117547.1-
Oryzias-melastigma':0.5172163305000002,(('XP_008403226.2-Poecilia-reticulata':0.10878663459999949,'XP_038162763.1-
Cyprinodon-tularosa':0.12897211139999953)'100/100':0.11200100220000042,'XP_037835926.1-Kryptolebias-
marmoratus':0.21187952859999992)'83.7/100':0.0336606159999997)'23/98':0.022644379500000866,'XP_041840514.1-
Melanotaenia-boesemani':0.18284952240000063)'100/100':0.07116668719999986,('XP_039454927.1-Oreochromis-
aureus':0.1952852237,'XP_035800389.1-Amphiprion-
ocellaris':0.17938009130000054)'18.6/81':0.02325029330000028)'90.3/100':0.01595186790000014,'XP_028999542.1-Betta-
splendens':0.29853974890000057)'77.6/88':0.015546731299999728,(('XP_020479276.1-Monopterus-
albus':0.1567714785000005,'XP_026149125.1-Mastacembelus-
armatus':0.10788492729999977)'98.8/98':0.02264639990000017,('XP_029313126.1-Cottoperca-
gobio':0.10835386739999997,'XP_034413570.1-Cyclopterus-
lumpus':0.08484388080000027)'99.9/100':0.03276545049999946)'93.1/88':0.012943654799999926)'91.6/99':0.0190922207000001
62)'98.7/79':0.03313830979999999,'XP_029688116.1-Takifugu-
rubripes':0.31556409969999955)'100/83':0.09406232370000023,'XP_030232486.1-Gadus-
morhua':0.3729581364000003)'96.6/83':0.04128118589999996,'XP_046897336.1-Hypomesus-
transpacificus':0.1974643533)'86.9/83':0.03406287149999976,'XP_042174965.1-Oncorhynchus-
tshawytscha':0.21355913449999964)'100/83':0.13818377590000086)'100/83':0.08885425989999973,'XP_035274752.1-Anguilla-
anguilla':0.1875541867999999)'62.9/82':0.03046621539999972,'XP_018618641.1-Scleropages-
formosus':0.19415536059999994)'100/83':0.1374535065,'XP_015198814.1-Lepisosteus-
oculatus':0.15205238799999954)'100/82':0.10777383220000036,'XP_034757766.1-Acipenser-
ruthenus':0.1706774962000006)'98.6/82':0.06067951090000001)'100/71':0.1442244799000001)'19.2/40':0.08567035280000024,('
XP_007905552.2-Callorhinchus-milii':0.2322661064000009,('XP_032906424.1-Amblyraja-
radiata':0.3595445937999999,'XP_048418965.1-Stegostoma-
fasciatum':0.22244403749999986)'100/100':0.16636698069999944)'98.3/100':0.14166330840000008)'100/100':1.971191524)'65.6/
40':0.1544286222000011)'97.1/76':0.16467188059999938)'100/100':0.6757021576)'19.6/82':0.13697024270000036)'43.9/93':0.196
7684794000002):0.7249423630000003);

#### Just ROC, COR-A, and kinase domains (tree shown in Figure S6).

Sequences: 273
Alignment length: 1099
Best fit model: Q.insect+I+G4

(((('XP_001134523.1-Dictyostelium-discoideum-AX4':0.08604597149999993,'XP_003284809.1-Dictyostelium-
purpureum':0.10722562980000028)'99.8/100':0.1655928793000001,('XP_012760012.1-Acytostelium-subglobosum-
LB1':0.1396003411000004,('XP_020428079.1-Heterostelium-album-PN500':0.0942905823000002,'XP_012749112.1-Acytostelium-
subglobosum-
LB1':0.2438108928)'45.4/94':0.04613939069999616)'99.5/100':0.1386093629999996)'100/100':0.7468195387999996,((((('XP_003
294476.1-Dictyostelium-purpureum':0.053158815799999815,'XP_645923.1-Dictyostelium-discoideum-
AX4':0.04973990019999963)'100/100':0.20897125659999993,'XP_004361995.1-Cavenderia-
fasciculata':0.15691418579999983)'50.3/98':0.04722042189999964,'XP_012757658.1-Acytostelium-subglobosum-
LB1':0.14278744609999983)'65.3/99':0.03141154440000005,'XP_020429574.1-Heterostelium-album-
PN500':0.06170227149999974)'100/100':0.5394359477000004,('XP_004352522.1-Acanthamoeba-castellanii-str.-
Neff':0.7341111627999997,('XP_004353012.1-Acanthamoeba-castellanii-str.-Neff':0.7387164063,(('XP_004337072.1-
Acanthamoeba-castellanii-str.-Neff':0.1634822667,'XP_004339596.1-Acanthamoeba-castellanii-str.-
Neff':0.2951896933000002)'99.3/100':0.17378499939999958,('XP_004367915.1-Acanthamoeba-castellanii-str.-
Neff':0.2588399061000004,'XP_004367596.1-Acanthamoeba-castellanii-str.-
Neff':0.18550644819999995)'98.7/100':0.14406366189999975)'100/100':0.32125454110000007)'97.1/100':0.16639004570000004)'
97.4/100':0.18960511419999992)'98.5/100':0.26970949970000024)'100/100':0.49367964135,(((((('XP_032810325.1-Petromyzon-
marinus':0.44262261639999956,(((('XP_041113508.1-Polyodon-spathula':0.10100687560000043,((('XP_018595881.2-
Scleropages-formosus':0.1710697211000003,'XP_048825230.1-Brienomyrus-
brachyistius':0.16600128690000027)'99.4/100':0.0712390433000003,(((('XP_041966047.1-Alosa-
sapidissima':0.16527686870000036,'XP_028813822.1-Denticeps-
clupeoides':0.1639050066000003)'99.4/100':0.06124987080000022,('NP_001188385.2-Danio-
rerio':0.24797382760000009,('XP_047675646.1-Tachysurus-fulvidraco':0.17464941079999985,('XP_026859030.2-Electrophorus-
electricus':0.1654047219999999,'XP_036431880.1-Colossoma-
macropomum':0.08746831190000037)'59/100':0.013571752999999909)'99.5/100':0.04989987739999968)'98.9/100':0.0460552378
0000009)'26.9/91':0.015074271599999634,'XP_030646389.1-Chanos-
chanos':0.26445595239999964)'92.6/100':0.02955418480000027,(('XP_030221087.1-Gadus-
morhua':0.27279227939999995,((((('XP_033824647.1-Periophthalmus-magnuspinnatus':0.27153891309999967,'XP_037110961.1-
Syngnathus-acus':0.2019620002)'90.8/99':0.037037072599999554,('XP_034390118.1-Cyclopterus-
lumpus':0.06312615830000023,'XP_037306590.1-Pungitius-
pungitius':0.08672125990000001)'66.1/99':0.019651132499999946)'91.8/87':0.021925072700000214,((('XP_028304639.1-
Gouania-willdenowi':0.14263941290000037,'XP_029951512.1-Salarias-
fasciatus':0.10731375130000043)'84.3/100':0.014368034500000348,'XP_005731602.1-Pundamilia-
nyererei':0.09744524450000025)'63.9/97':0.009628172300000237,(('XP_015819282.1-Nothobranchius-
furzeri':0.10402986049999985,('XP_037834279.1-Kryptolebias-marmoratus':0.07730353549999958,'XP_013876060.1-
Austrofundulus-
limnaeus':0.10276193829999958)'98.2/100':0.03254806809999966)'1.1/57':0.02152729380000018,'XP_024137746.1-Oryzias-
melastigma':0.17732088160000004)'97.8/99':0.027780536599999905)'68.3/94':0.00445684540000002)'61.3/85':0.0093120861999
9957,('XP_029010427.1-Betta-splendens':0.14872217080000016,'XP_026220797.1-Anabas-
testudineus':0.047295765400000356)'97.4/99':0.03484754599999995)'97.2/99':0.029424394600000348,'XP_046906764.1-
Hypomesus-
transpacificus':0.16558032509999965)'44.1/82':0.013109925000000189)'66.1/88':0.015553414500000251,('XP_010896173.2-
Esox-lucius':0.08830567469999995,'XP_024235857.1-Oncorhynchus-
tshawytscha':0.05109107360000314)'98.5/100':0.03943686339999744)'97.4/100':0.04272468170000021)'99.7/100':0.063532402
39999991)'88.6/100':0.021164399699999947,('XP_035281064.1-Anguilla-anguilla':0.11397914369999995,'XP_036373397.1-
Megalops-
cyprinoides':0.15829785509999983)'98.4/100':0.04399689300000009)'100/100':0.1949360975000003)'33/86':0.022070281000000
413,'XP_039617043.1-Polypterus-senegalus':0.15638291329999987)'100/100':0.1474015111,(((('XP_044136143.1-Bufo-
gargarizans':0.11308131270000032,'XP_018108120.1-Xenopus-
laevis':0.1010887198999999)'99.8/100':0.0719085089,'XP_029473217.1-Rhinatrema-
bivittatum':0.13714243250000013)'35.5/92':0.01632003419999961,(((((((('XP_037369231.1-Talpa-
occidentalis':0.27905191859999956,'XP_016047443.1-Erinaceus-
europaeus':0.05378772909999974)'99.5/100':0.04654991459999991,'XP_049639831.1-Suncus-
etruscus':0.04510806079999963)'91.5/100':0.01300793510000009,'XP_046527794.1-Equus-
quagga':0.011112909899999579)'84/84':0.004447525999999868,((('XP_027253651.1-Cricetulus-
griseus':0.07862015300000014,'XP_040609888.1-Mesocricetus-
auratus':0.012433894099999954)'93.6/100':0.012188832199999666,'XP_036063763.1-Onychomys-
torridus':0.015933015899999958)'66/100':0.002691021899995905,'XP_049987333.1-Microtus-
fortis':0.02372118450000027)'98.4/100':0.02000986440000041)'81.2/81':0.004314948300000232,'NP_940980.4-Homo-sapiens-
LRRK2':0.0239565715000003)'47.7/80':0.006224639900000106,'XP_023564370.1-Octodon-
degus':0.0374270656000002)'99.9/100':0.05821671340000023,'XP_043824509.1-Dromiciops-
gliroides':0.06158890170000042)'99.7/100':0.044650721000000004,((((('XP_032077596.1-Thamnophis-
elegans':0.01814029240000035,'XP_026533023.1-Notechis-
scutatus':0.04751028040000005)'100/100':0.1411568113000013,'XP_020652645.1-Pogona-
vitticeps':0.0361745673999998)'60.3/100':0.010638457599999818,'XP_034982363.1-Zootoca-

vivipara':0.08419830670000028)'75.9/99':0.00876207099999594,'XP_044279156.1-Varanus-
komodoensis':0.05845704610000002)'99.7/100':0.03641308440000035,('XP_034622773.1-Trachemys-scripta-
elegans':0.0569567150000001,('XP_025065501.1-Alligator-sinensis':0.03557916809999995,('XP_009555590.1-Cuculus-
canorus':0.013190438899999712,('XP_030816227.1-Camarhynchus-parvulus':0.01461001779999993,'XP_017666944.1-
Lepidothrix-
coronata':0.008976696899999581)'85.4/100':0.007223972800000311)'100/100':0.04922143830000003)'97.8/100':0.030779035000
00012)'46.5/99':0.009569948300000242)'97.3/100':0.022034431899999873)'99.5/100':0.04360725709999686)'99.7/100':0.059486
473999999845,'XP_014347482.1-Latimeria-
chalumnae':0.10056890220000003)'95.6/100':0.048221837399999856)'97.5/100':0.0941317922999998,('XP_007908179.2-
Callorhinchus-milii':0.08403323060000023,('XP_038668138.1-Scyliorhinus-canicula':0.05412214089999967,('XP_048410179.1-
Stegostoma-fasciatum':0.02862269580000021,'XP_043569546.1-Chiloscyllium-
plagiosum':0.05169006369999973)'99/100':0.03982416769999997)'100/100':0.09369974240000012)'81.8/100':0.06642219759999
968)'100/100':0.3959015854999999)'100/100':0.47874558879999984,'XP_019623007.1-Branchiostoma-
belcheri':0.2953514394000001)'79.5/86':0.0640587198999978,(('XP_029643290.1-Octopus-
sinensis':0.8430944330000001,(('XP_041348340.1-Gigantopelta-aegis':0.4317774379000001,'XP_048252841.1-Haliotis-
rufescens':0.23167786920000033)'5.3/57':0.04628789809999967,('XP_035828109.1-Aplysia-
californica':0.20486370139999988,'XP_025107350.1-Pomacea-
canaliculata':0.24952036430000035)'100/100':0.13617604829999985)'99.4/100':0.11259637160000002)'70.9/98':0.034437583799
99995,('XP_033759237.1-Pecten-maximus':0.4052003420000001,('XP_034308389.1-Crassostrea-
gigas':0.08630960059999992,'XP_022298272.1-Crassostrea-
virginica':0.05607799629999999)'100/100':0.28880428349999976)'99.4/100':0.14356825989999988)'100/100':0.276095128200000
15)'85.1/86':0.080501092,('XP_030828896.1-Strongylocentrotus-purpuratus':0.3171883507000004,('XP_033645351.1-Asterias-
rubens':0.06665976520000028,('XP_038069546.1-Patiria-miniata':0.036491387599999925,'XP_022097750.1-Acanthaster-
planci':0.03605327699999972)'99.2/100':0.06408998679999955)'100/100':0.20708843850000003)'100/100':0.2741193675)'87.4/93'
:0.09098127820000013,('XP_028391624.1-Dendronephthya-gigantea':1.0795014026000005,(('XP_022785899.1-Stylophora-
pistillata':0.26005247129999987,'XP_020610026.1-Orbicella-
faveolata':0.11157646749999994)'100/100':0.3398103146999998,('XP_048584778.1-Nematostella-
vectensis':0.2003167925999998,('XP_031554299.1-Actinia-tenebrosa':0.19720493139999995,'XP_020898149.1-Exaiptasia-
diaphana':0.32614598269999995)'96.6/100':0.09626904859999996)'99.5/100':0.16357442399999966)'98.9/100':0.2081346630000
0005)'99.8/100':0.3041422645999998)'99.9/100':0.46527656509999993,((((((('XP_035701600.1-Folsomia-
candida':0.5772887704,('XP_003373529.1-Trichinella-spiralis':0.2802279960999998,('XP_042935833.1-Brugia-
malayi':0.2263168678999996,('NP_492839.4-Caenorhabditis-elegans':0.6485611976000003,'XP_024503827.1-Strongyloides-
ratti':0.8828516114999996)'97.9/100':0.17609609079999977)'99.1/100':0.12707383820000003)'99.9/100':0.1640128504999998)'63.
9/88':0.04984003050000041,((('XP_027206363.1-Dermatophagoides-pteronyssinus':0.0241473430000001,'XP_046912895.1-
Dermatophagoides-farinae':0.0284980047000003)'100/100':0.49103999819999977,'XP_015784227.1-Tetranychus-
urticae':0.18009138179999962)'93.2/90':0.04663569089999964,((((((((((((('XP_013174285.1-Papilio-
xuthus':0.10875549619999969,('XP_049883152.1-Pectinophora-gossypiella':0.1304321169999998,(('XP_037875629.1-Bombyx-
mori':0.12134258539999987,'XP_037299202.1-Manduca-
sexta':0.09659393050000009)'94.7/96':0.026127214499999774,'XP_026747625.1-Trichoplusia-
ni':0.07803077179999995)'92.8/97':0.0258515905000003)'97.6/100':0.0397616765000004)'72.8/99':0.02179576700000041,(('XP_0
45530211.1-Pieris-brassicae':0.10517610560000001,'XP_041989149.1-Aricia-
agestis':0.11117533230000021)'82.4/98':0.0156824677999996,(('XP_032525892.1-Danaus-plexippus-
plexippus':0.12706472200000007,('XP_047542335.1-Vanessa-atalanta':0.011109461300000234,'XP_046975207.1-Vanessa-
cardui':0.007615873499998146)'100/100':0.05146991929999967)'37.6/97':0.013214434100000005,'XP_023937942.1-Bicyclus-
anynana':0.061628690400000075)'98.4/99':0.03167014200000029)'95.8/98':0.03662124249999987)'100/100':0.5138318087,(((('X
P_049531803.1-Anopheles-darlingi':0.008548864499999809,('XP_050083787.1-Anopheles-aquasalis':0.000003,'XP_035783724.1-
Anopheles-
albimanus':0.01171587289999998)'43.8/100':0.003619047199999474)'100/100':0.07006693900000016,'XP_050071280.1-
Anopheles-maculipalpis':0.031577995000000136)'98.7/100':0.03552719039999985,'XP_029736246.1-Aedes-
albopictus':0.02744035250000021)'99.8/100':0.07143559310000036,(('XP_017848820.1-Drosophila-
busckii':0.07407608120000031,'XP_032576550.1-Drosophila-
sechellia':0.04318812909999806)'100/100':0.08040946090000034,('XP_037942768.1-Teleopsis-
dalmanni':0.05268187300000046,(('XP_037891855.1-Glossina-fuscipes':0.04166694879999966,'XP_013097712.1-Stomoxys-
calcitrans':0.02448916049999994)'56.2/100':0.007224794099999876,('XP_028897545.1-Zeugodacus-
cucurbitae':0.04336848029999845,'XP_020713747.1-Ceratitis-
capitata':0.0769571971999996)'100/100':0.0500025559999992)'39.8/89':0.006835101500000107)'91.5/89':0.0199384248000003)'
100/100':0.2188619062999999)'97.7/100':0.0704630790999996)'99.7/100':0.10647825489999985,((((('XP_014209526.1-
Copidosoma-floridanum':0.06828729949999968,'XP_014233075.1-Trichogramma-
pretiosum':0.08851911659999967)'20.8/94':0.009938370699999588,'XP_003425729.1-Nasonia-
vitripennis':0.01625793799999986)'99.8/96':0.03452140240000234,(('XP_017885653.1-Ceratina-
calcarata':0.08747687959999961,'XP_043263337.1-Colletes-
gigas':0.0630521813999998)'71.9/99':0.01142140580000173,'XP_026829906.1-Ooceraea-
biroi':0.0464471611999997)'95.8/98':0.01718871130000215)'82.9/78':0.00672032960000241,(('XP_044005005.1-Aphidius-
gifuensis':0.10840524099999982,'XP_015116519.1-Diachasma-
alloeum':0.04274420700000145)'92.7/98':0.02224570020000005,'XP_043282449.1-Venturia-
canescens':0.04341002329999988)'30.8/71':0.004154364299999713)'75.6/50':0.013038270200000035,'XP_024943937.1-Cephus-
cinctus':0.02603517990000004)'100/100':0.1952966010999999)'90.6/92':0.03194856719999972,('XP_044740580.1-Chrysoperla-
carnea':0.05898772559999976,((('XP_045460940.1-Harmonia-axyridis':0.18633602299999996,'XP_015840646.1-Tribolium-
castaneum':0.02213749309999794)'15.3/91':0.006997307399999819,('XP_048526705.1-Dendroctonus-
ponderosae':0.06971758400000017,'XP_023029002.1-Leptinotarsa-

decemlineata':0.0387195801000004 3)'87.5/100':0.01366271499999714)'87.7/97':0.00990786960000119,(('XP_018323263.1-
Agrilus-planipennis':0.06498711470000007,'XP_031347202.1-Photinus-
pyralis':0.04890933849999968)'96.7/100':0.02701193209999747,'XP_017772148.1-Nicrophorus-
vespilloides':0.03338609770000023)'64.6/99':0.01391566349999973)'100/100':0.05237712649999704)'70.3/99':0.0134140758000
00091)'88.4/88':0.02437806549999657,('XP_002427797.1-Pediculus-humanus-corporis':0.2306961536000003,'XP_026273351.1-
Frankliniella-
occidentalis':0.1180291270999998)'86.6/90':0.03242184229999978)'87.4/88':0.01440456410000035,('XP_021920935.1-
Zootermopsis-nevadensis':0.04262899560000033,'XP_049788729.1-Schistocerca-
nitens':0.04824052449999715)'90.4/100':0.01518389369999729)'94.9/88':0.02706380089999713,((('XP_025421547.1-Sipha-
flava':0.2191908030000005,('XP_014276941.1-Halyomorpha-halys':0.06303838619999969,'XP_014247457.1-Cimex-
lectularius':0.05960561920000007)'100/100':0.1677982216000002)'34/62':0.02394503649999794,'XP_018906419.1-Bemisia-
tabaci':0.12323639929999963)'3.3/24':0.01587340689999942,'XP_046668017.1-Homalodisca-
vitripennis':0.05900181179999997)'96.2/92':0.02633606679999989)'98.7/97':0.03788459440000036,'XP_046393381.1-Ischnura-
elegans':0.09728150030000027)'100/100':0.0827951879000004,'XP_045599192.1-Procambarus-
clarkii':0.1543076037000004)'93.5/97':0.03240560740000314,'XP_043233053.1-Amphibalanus-
amphitrite':0.20368568479999993)'52.2/74':0.01348120050000234,(('XP_040578378.1-Lepeophtheirus-
salmonis':0.3598219160999996,'XP_023326413.1-Eurytemora-
affinis':0.18928739299999986)'100/100':0.13654707450000014,'XP_046646629.1-Daphnia-
pulicaria':0.24094849569999965)'77.4/67':0.03427626219999685)'99.9/95':0.07127158100000042,'XP_042905583.1-
Parasteatoda-tepidariorum':0.1540073815999996)'78.7/77':0.027819668400000275,(('XP_028968531.1-Galendromus-
occidentalis':0.053526703600000225,'XP_022703572.1-Varroa-
jacobsoni':0.05024514269999969)'100/100':0.24687129480000003,('XP_042147598.1-Ixodes-
scapularis':0.07670865079999967,'XP_049528486.1-Dermacentor-
silvarum':0.16548835429999986)'99.7/100':0.07717055110000004)'97.3/100':0.0515391199000015)'96.9/78':0.040538971200000
16)'93.4/90':0.04517349049999986)'100/100':0.1572396896999999,(('XP_029646318.1-Octopus-
sinensis':0.000691,'XP_014782789.1-Octopus-bimaculoides':0.006211990299999748)'100/100':0.5118182915,(('XP_045213481.1-
Mercenaria-mercenaria':0.6159767617999998,('XP_033739213.1-Pecten-maximus':0.3121811658000002,('XP_048762622.1-
Ostrea-edulis':0.08211853740000041,'XP_034314769.1-Crassostrea-
gigas':0.04998121169999584)'100/100':0.23390997040000006)'100/100':0.21349544470000037)'23.7/92':0.08311309359999974,
('XP_013066303.1-Biomphalaria-glabrata':0.41101563009999964,('XP_041363499.1-Gigantopelta-
aegis':0.3264924875000004,'XP_048248699.1-Haliotis-
rufescens':0.19790442670000008)'65.6/99':0.07623303819999983)'99.7/100':0.1631243396000004)'98.4/100':0.115692537600000
2)'100/100':0.42642412100000016)'96.8/97':0.09944645099999994,('XP_019617338.1-Branchiostoma-
belcheri':0.5062968482000003,('XP_033120526.1-Anneissia-japonica':0.16431012199999984,('XP_041462892.1-Lytechinus-
variegatus':0.16704773469999967,('XP_022107621.1-Acanthaster-planci':0.07337287589999963,'XP_033635275.1-Asterias-
rubens':0.1065645743000001)'100/100':0.13852743410000024)'100/100':0.1346431680000002)'100/100':0.3834756962000001)'54
.1/96':0.07222850660000013)'99.1/97':0.18669372969999998,(('XP_047128450.1-Hydra-
vulgaris':0.2455376208000004,'XP_047128453.1-Hydra-
vulgaris':0.13984886680000042)'100/100':0.7971456382000004,('XP_028415446.1-Dendronephthya-
gigantea':0.4995497830000004,(('XP_022795822.1-Stylophora-pistillata':0.08182386529999963,'XP_029187052.2-Acropora-
millepora':0.10573835739999993)'100/100':0.11941623540000013,('XP_031568177.1-Actinia-
tenebrosa':0.08554520339999971,'XP_048577767.1-Nematostella-
vectensis':0.09702996830000021)'99.4/100':0.10884658989999973)'99.8/100':0.1732589657000001)'99.3/100':0.18323298530000
01)'95.1/100':0.1370854843)'100/100':0.6991792482000001,(('XP_012555367.2-Hydra-
vulgaris':0.6465172126000001,('XP_047143158.1-Hydra-vulgaris':0.2681784186999998,('XP_047143281.1-Hydra-
vulgaris':0.17480058289999967,(('XP_047144213.1-Hydra-vulgaris':0.06311144489999965,'XP_047144101.1-Hydra-
vulgaris':0.08935861780000032)'98.9/100':0.05815552949999958,'XP_047143514.1-Hydra-
vulgaris':0.08417560909999988)'93.7/100':0.0654049527999998)'94.7/99':0.11036861559999966)'99.6/100':0.2855766211999997)
'100/100':0.8766578197999997,('XP_031569514.1-Actinia-tenebrosa':1.3073516211999996,(('XP_028409574.1-Dendronephthya-
gigantea':0.29412121629999977,'XP_046847823.1-Xenia-sp.-Carnegie-
2017':0.32960685259999956)'100/100':0.5230738266000001,('XP_032226651.2-Nematostella-
vectensis':0.19815584579999967,('XP_031571669.1-Actinia-tenebrosa':0.1670616278999999,'XP_020891986.1-Exaiptasia-
diaphana':0.18310764499999976)'94.2/100':0.12355859579999962)'100/100':0.48937645350000025)'91.1/100':0.17227368090000
006)'87.4/99':0.1433614846000002)'100/100':0.47780460280000003)'53.5/81':0.08117111010000011,(('XP_047142009.1-Hydra-
vulgaris':0.7132694898,(('XP_046856792.1-Xenia-sp.-Carnegie-2017':0.16069705429999992,'XP_028416799.1-Dendronephthya-
gigantea':0.14877950049999988)'100/100':0.4434301507999998,(('XP_015772409.1-Acropora-
digitifera':0.1705404118999998,('XP_020612099.1-Orbicella-faveolata':0.06136668710000004,'XP_027055244.1-Pocillopora-
damicornis':0.0972735076000002)'73.7/100':0.03355666180000054)'100/100':0.18289326189999988,('XP_048583427.1-
Nematostella-vectensis':0.2806958987999999,('XP_020914476.1-Exaiptasia-diaphana':0.15785067330000002,'XP_031560456.1-
Actinia-
tenebrosa':0.10990703190000017)'97/100':0.06646661810000021)'98.6/100':0.12208609649999991)'100/100':0.27080961650000
024)'87.1/99':0.11359095880000014)'99.5/100':0.1998010111999993,(('XP_045189912.1-Mercenaria-
mercenaria':0.8407294952000002,(('XP_022329885.1-Crassostrea-virginica':0.1366229647999999,'XP_048737827.1-Ostrea-
edulis':0.06767462230000021)'100/100':0.502153946,('XP_021368573.1-Mizuhopecten-
yessoensis':0.16051860809999985,'XP_033749578.1-Pecten-
maximus':0.09656691110000004)'100/100':0.5350199392000001)'29.4/94':0.09895797820000007)'100/100':0.6949470634999999,
('XP_002736404.1-Saccoglossus-kowalevskii':0.7869667347,((('XP_033109441.1-Anneissia-
japonica':1.1624071061999999,('XP_041464429.1-Lytechinus-variegatus':0.8602642015000002,('XP_033633998.1-Asterias-
rubens':0.29204066000000006,(('XP_038074856.1-Patiria-miniata':0.1373888859000001,'XP_038074850.1-Patiria-
miniata':0.11286429570000012)'100/100':0.22601791439999985,('XP_022111022.1-Acanthaster-

planci':0.12176772049999984,'XP_038067103.1-Patiria-
miniata':0.1539131134999998)'26.5/98':0.04199244369999988)'99.7/99':0.1841468411)'100/100':0.40061031830000005)'99.9/100'
:0.23900433480000016)'9.8/75':0.07223590909999977,'XP_033111272.1-Anneissia-
japonica':0.9880993612999998)'99.7/100':0.26693804389999976,((((('XP_040198253.1-Rana-
temporaria':0.0889840552999992,'XP_041442810.1-Xenopus-
laevis':0.06987939909999952)'100/100':0.09770456790000015,'XP_030045605.1-Microcaecilia-
unicolor':0.1051913175000001)'96.5/100':0.03386450670000052,((((('XP_024897004.1-Pteropus-
alecto':0.11859348189999963,('XP_014399058.1-Myotis-brandtii':0.11972098790000052,'XP_036211673.1-Myotis-
myotis':0.00386030930000253)'99.6/100':0.03677126269999942)'66.9/74':0.00611391179999983,('XP_012588204.1-Condylura-
cristata':0.02747535149999969,'NP_078928.3-Homo-sapiens-
LRRK1':0.01790331639999998)'52.8/97':0.005063185700000835)'84.6/78':0.00472631970000481,('XP_042844148.1-Panthera-
tigris':0.00652406309999644,'XP_032478864.1-Phocoena-
sinus':0.02333571640000095)'74.3/88':0.00252258700000052)'91.8/97':0.01316888000000271,'XP_045142310.1-Echinops-
telfairi':0.06389598860000056)'100/100':0.06342372279999964,((('XP_032088514.1-Thamnophis-
elegans':0.02457786839999758,'XP_039215026.1-Crotalus-
tigris':0.02719296220000004)'100/100':0.08555732810000016,('XP_048338174.1-Sphaerodactylus-
townsendi':0.08807547329999998,'XP_034986799.1-Zootoca-
vivipara':0.05234385149999987)'91.2/100':0.01217888780000204)'99.9/100':0.03576446589999982,('XP_019333123.1-Alligator-
mississippiensis':0.03415666639999948,'XP_025977499.1-Dromaius-
novaehollandiae':0.04597449809999965)'81.5/97':0.00571605569999778)'85.5/97':0.02180316180000304)'97.6/100':0.0345055
3610000018)'100/100':0.0895134809,('XP_039628634.1-Polypterus-senegalus':0.13554845609999955,((((((((('XP_016103532.1-
Sinocyclocheilus-grahami':0.01853341720000457,'XP_026071877.1-Carassius-
auratus':0.03904013130000017)'100/100':0.08811332979999964,'XP_046713878.1-Silurus-
meridionalis':0.08793077749999956)'22.2/59':0.02177690829999675,'XP_035387696.1-Electrophorus-
electricus':0.07740107179999978)'98.4/100':0.0425349430000006,('XP_028812427.1-Denticeps-
clupeoides':0.10868530079999994,'XP_041917165.1-Alosa-
sapidissima':0.06355712850000028)'45.2/69':0.01434134820000326)'92.6/98':0.02064897039999991,((((((('XP_033847562.1-
Periophthalmus-magnuspinnatus':0.2632835588,'XP_029977649.1-Sphaeramia-
orbicularis':0.07668377420000017)'73.7/80':0.02272003180000004,'XP_041640649.1-Cheilinus-
undulatus':0.09721157380000012)'90.2/49':0.01062613390000223,('XP_028333065.1-Gouania-
willdenowi':0.15824780010000028,('XP_008308656.1-Cynoglossus-semilaevis':0.10405883490000001,'XP_043886127.1-Solea-
senegalensis':0.11816649540000013)'96/78':0.0330063827)'71.7/47':0.01106429460000135)'88.3/44':0.00626807379999829,(((('
XP_029688116.1-Takifugu-rubripes':0.14302605320000028,'XP_034413570.1-Cyclopterus-
lumpus':0.03297580460000038)'90.4/82':0.01108883179999532,'XP_029313126.1-Cottoperca-
gobio':0.05032612549999982)'81.2/76':0.00976978080000127,('XP_020479276.1-Monopterus-
albus':0.06707423250000044,'XP_026149125.1-Mastacembelus-
armatus':0.0576500029)'94.6/100':0.01524455589999845)'89.4/64':0.00825902849999932)'65.4/54':0.00394185829999712,(((('
XP_024117547.1-Oryzias-melastigma':0.2879006035999998,'XP_041840514.1-Melanotaenia-
boesemani':0.06119904259999975)'77.9/98':0.01996982680000146,(('XP_008403226.2-Poecilia-
reticulata':0.0567963565999996,'XP_038162763.1-Cyprinodon-
tularosa':0.08870677120000003)'100/100':0.05886781890000048,'XP_037835926.1-Kryptolebias-
marmoratus':0.09994541760000031)'79.8/100':0.01694231009999969)'98.8/98':0.03278652920000025,('XP_039454927.1-
Oreochromis-aureus':0.09249358059999979,'XP_035800389.1-Amphiprion-
ocellaris':0.09608423949999967)'24/58':0.01122684940000294)'90.9/75':0.01079916500000333,'XP_028999542.1-Betta-
splendens':0.14620358990000026)'46.8/54':0.00466352219999548)'100/85':0.04983775410000035,('XP_030232486.1-Gadus-
morhua':0.1513456743999999,'XP_046897336.1-Hypomesus-
transpacificus':0.0866633948000004)'40/92':0.01261572760000007)'92/85':0.01422322019999886,'XP_042174965.1-
Oncorhynchus-
tshawytscha':0.06767381439999998)'100/85':0.06468200860000017)'98/84':0.03836407879999992,'XP_035274752.1-Anguilla-
anguilla':0.08197614210000026)'46.7/83':0.01691058920000103,'XP_018618641.1-Scleropages-
formosus':0.09283018400000032)'99.5/83':0.04895752629999972,'XP_015198814.1-Lepisosteus-
oculatus':0.05802267500000135)'99.1/83':0.04281943769999952,'XP_034757766.1-Acipenser-
ruthenus':0.08724676979999924)'99.1/83':0.04240818550000025)'98.5/79':0.05656927790000436)'86/78':0.06081186149999951
,('XP_007905552.2-Callorhinchus-milii':0.1228898052000007,('XP_032906424.1-Amblyraja-
radiata':0.2030567739000002,'XP_048418965.1-Stegostoma-
fasciatum':0.11367109640000006)'94.5/92':0.06007109619999973)'39.8/74':0.02844293850000046)'100/100':1.093839263999999
6)'71.8/92':0.10464903220000021)'81.1/88':0.07499420870000018)'82.8/92':0.09226121719999991)'100/100':0.296124146399999
95)'89.3/87':0.17070371299999998):0.49367964135);
